# Analysis of blood-based DNA Methylation signatures of aging and disease progression in IBD patients

**DOI:** 10.1101/2025.02.08.635564

**Authors:** Trevor Doherty, Edel McDermott, Sarah Jane Delany, Hugh Mulcahy, Therese M. Murphy

## Abstract

**Background:** Inflammatory bowel diseases (IBDs) are chronic inflammatory disorders with a dysregulated immune response partly influenced by environmental factors. DNA methylation (DNAm), a key epigenetic mechanism, is implicated in the etiology of complex diseases, including IBD. Epigenetic clocks, which use DNAm patterns to estimate biological aging, have been increasingly linked to various health and disease states. Previous studies have associated DNAm with IBD, and first- and second-generation epigenetic clocks with IBD subtypes.

**Results:** In a discovery IBD cohort (n=149) with 8-year clinical follow-up data, we explored the relationship between DNAm variation, second- and third-generation epigenetic clocks, and IBD clinicopathological outcomes, including disease subtype, activity, and recurrence. One CpG site was significantly differentially methylated (Benjamini-Hochberg adjusted p-value<0.05) in patients with clinical recurrence of disease over the long term (i.e., after the first year of study) compared to non-recurrence (no treatment escalation after 8 years). Next, we assessed DNAm aging signatures and IBD outcomes using logistic regression. Individuals with IBD exhibited significantly increased epigenetic aging, as measured by GrimAge, GrimAge2, and DunedinPACE, compared with controls. These associations were replicated in two independent IBD cohorts (GSE87648 (n=377) and GSE112611 (n=238)). Additionally, in UC patients, the active disease group was associated with higher age acceleration (GrimAge (U=669, p=0.003)) and higher pace of aging (DunedinPACE (t=3.233, 0.002)) compared to the inactive group. In the discovery cohort, DunedinPACE outperforms CRP measures in discriminating activity in UC patients with an AUC, sensitivity and specificity of 0.71, 69.5% and 68.7% respectively, highlighting its potential as a useful biomarker of activity in UC.

**Conclusions:** Overall, we present strong evidence that dynamic age-related DNAm changes can be used to differentiate between IBD (including separately by subtype) and controls. Furthermore, our study provides important new evidence that DunedinPACE may have utility as a biomarker for monitoring disease recurrence in IBD patients and may be a strong marker of disease activity in UC patients. Overall, this suggests that blood-based DNAm signatures could serve as biomarkers for detection and monitoring of IBD.

## Background

Inflammatory bowel diseases (IBDs), which encompass Crohn’s disease (CD) and ulcerative colitis (UC), are chronic inflammatory disorders impacting the gastrointestinal tract, characterised by alternating periods of flare-ups and remissions. These conditions can progress and lead to complications such as abscesses, colitis-associated neoplasia, and cancer (1). The etiology of IBD involves a complex interplay among genetic variation, the gut microbiota, the immune system, and environmental factors. Although genome-wide association studies (GWAS) have identified over 215 genetic loci associated with IBD susceptibility (2), only a fraction of the disease’s variability can be explained by genetic factors, highlighting the significant contribution of non-genetic elements (3). Current research increasingly investigates epigenetic mechanisms, such as DNA methylation (DNAm), which can regulate gene expression through modifications to DNA, histone proteins, and chromatin. DNAm changes have been implicated in various conditions, including cancer (4), aging (5–8) and IBD (9). Studies indicate that disease-associated DNAm variations observed in peripheral tissues may reflect patterns in colonic mucosa (3, 10, 11).

Presently, gastroenterologists rely on established clinical predictors and immunological/serological biomarkers to guide management, but conflicting reports exist regarding their prognostic value (12). Thus, accurate prognostication at IBD diagnosis remains a crucial research priority (13). Existing studies on DNAm changes in IBD are predominantly cross-sectional, with limited exploration of the predictive value of epigenetic-based predictors for monitoring disease activity and recurrence (9, 14, 15).

Recent studies have identified a significant correlation between epigenetic age acceleration (defined as the discrepancy between predicted age, as determined by DNAm patterns, and chronological age) and IBD, as well as its subtypes (15, 16). The development of second-generation epigenetic clocks, such as the PhenoAge (17) and GrimAge clocks (7, 18), and the third-generation DunedinPACE (8) clock represents a notable advancement in this field. These clocks are trained to estimate aging-associated phenotypes and diseases, either instead of or in addition to chronological age, and have demonstrated superior performance compared to their predecessors. They offer enhanced accuracy in associating with chronic diseases and predicting expected lifespan. However, the application and evaluation of next generation epigenetic clocks in relation to IBD, its subtypes, and disease progression remain largely unexplored. Investigating these advanced epigenetic clocks could provide deeper insights into the molecular mechanisms underpinning IBD and potentially lead to more precise prognostic tools and therapeutic strategies.

Previously, we conducted DNA methylation profiling in peripheral blood from adult IBD patients and controls using the Illumina 450K microarray (3). Here, we expand on this dataset with new 8-year clinical follow-up data from the original IBD patient cohort (n=149), including relevant clinical information relating to disease progression and treatment response collected. In this study, we examine DNA methylation signatures in the original cohort and their association with disease recurrence, activity and subtypes in IBD patients.

## Methods

### Study Demographics

Three datasets were used in the analyses. These comprised a discovery dataset (3) and two replication datasets sourced from the Gene Expression Omnibus (GSE112611 and GSE87648). In each case, DNAm was quantified from whole blood samples. Further details of each dataset and study are provided below.

### Discovery Cohort

The discovery cohort (IBD cases (n=149), controls (n=39)) used in our analysis was previously enrolled in a study (3) while attending a university teaching hospital between July 2011 and November 2013. Eight-year clinical follow-up of this patient population was completed with relevant clinical information relating to disease progression and treatment response collected. Our primary clinical endpoint for our discovery cohort is clinical recurrence of disease activity requiring treatment escalation (19, 20) over short (treatment escalation in the first study year, anticipating that baseline active disease would largely be treated over this time period (21) and long term (after the first year of study (22)). Treatment escalation (recorded as 1 (yes)/0 (no)) was defined as the need for additional medical (new steroid, immunomodulator or biologic) or surgical therapy in response to active disease, in accordance with national and international guidelines (13, 20, 22). Treatment changes relating to drug intolerance or in response to therapeutic drug monitoring without inflammatory activity, surgery without intestinal resection and prescription renewals following periods of noncompliance were not included as treatment escalation endpoints (20). Ethical approval was obtained from St. Vincent’s University Hospital (Dublin) Research and Ethics Committee. Recurrence information was not available for 3 IBD samples and age was missing for a single control. Therefore, the final analyses included 184 samples (median age 35 years, 97 male), comprising 146 IBD cases (87 CD and 59 UC) and 38 non-IBD controls, all of which had no history of inflammatory disorders. Further study population details have been reported previously (3). Details of the discovery cohort demographics are presented in Table 1. Additionally, details of disease recurrence and activity are outlined in Table 2 and Table 3 respectively.

**Table 1:** Discovery Cohort Demographics.

|  | IBD |  |  |
| --- | --- | --- | --- |
|  | CD | UC | Controls |
| **N** | 87 | 59 | 38 |
| **Age (median, IQR)** | 33 (25 – 40.5) | 41 (32 – 55.5) | 41 (28 - 52) |
| **Females, %** | 43.7% | 45.8% | 50.9% |
| **Smoking %**<br>**(Current/Former/Never)** | 47/23/17 | 29/24/6 | 23/9/6 |

**Table 2:**
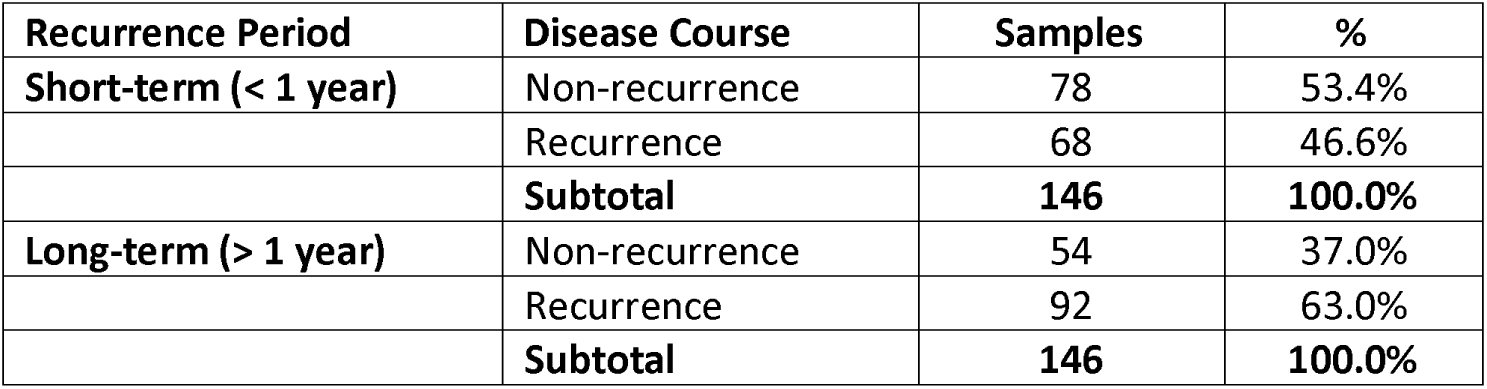
Discovery cohort – short and long-term recurrence status.

| Recurrence Period | Disease Course | Samples | % |
| --- | --- | --- | --- |
| **Short-term (< 1 year)** | Non-recurrence | 78 | 53.4% |
|  | Recurrence | 68 | 46.6% |
|  | **Subtotal** | **146** | **100.0%** |
| **Long-term (> 1 year)** | Non-recurrence | 54 | 37.0% |
|  | Recurrence | 92 | 63.0% |
|  | **Subtotal** | **146** | **100.0%** |

**Table 3:** Discovery Cohort – disease activity at baseline (active/inactive).

| Disease Activity | Samples | % |
| --- | --- | --- |
| **Inactive** | 70 | 47.9% |
| **Active** | 76 | 52.1% |
| **Total** | **146** | **100.0%** |

### Replication Cohorts

The first replication cohort (n=379, median age 33 years, 195 male) used in our analysis included patients recruited as part of the IBD-BIOM inception cohort and symptomatic controls recruited from gastroenterology clinics during the same period as described previously (23). The dataset was obtained from the online repository Gene Expression Omnibus (GEO) under the accession code GSE87648. Study demographic details are shown in Table 4.

**Table 4:** GSE87648 IBD Study Demographics.

|  | IBD | CD | UC | Controls |
| --- | --- | --- | --- | --- |
| **N** | 204 | 103 | 101 | 175 |
| **Age (median, IQR)** | 34 (25 - 49.3) | 32 (25 - 51) | 35 (27 - 47) | 32 (26 - 43) |
| **Females, %** | 46.6% | 48.5% | 44.6% | 50.9% |

The second replication cohort (n=238, median age 12.6, 136 male) involved paediatric subjects recruited under the Risk Stratification and Identification of Immunogenetic and Microbial Markers of Rapid Disease Progression in Children with Crohn’s Disease (RISK) study, as described previously (24). The study recruited children between ages 1 and 17 who presented to gastroenterology clinics with suspected IBD. The dataset was obtained from the GEO repository (accession code GSE112611). Details of the cohort demographics are presented in Table 5.

**Table 5:** GSE112611 Paediatric Study Demographics.

|  | CD | Controls |
| --- | --- | --- |
| **N** | 164 | 74 |
| **Age (median, IQR)** | 12.6 (10.6 - 15) | 12.6 (9.8 - 14.3) |
| **Females %** | 41% | 46% |

### DNA Methylation Microarray Pre-processing

Microarray preprocessing for the discovery cohort was carried out as described previously (3).

GEOquery (25) from the Bioconductor package in R was used to download both IDAT files and phenotype data from the GEO repository for the replication cohort GSE87648. DNAm was quantified using the Infinium HumanMethylation450 BeadChip (Illumina Inc., San Diego, CA) as described previously (23). For the epigenetic clock association analyses, the following quality control steps were implemented for the above GEO data set and the discovery cohort using the R Bioconductor minfi package (26). Samples where > 1% of sites had a corresponding detection p-value of > 0.01 were removed, as well as those with a bead count < 3 in 1% of samples. Similarly, CpGs where > 1% of samples had detection p-values > 0.01 were removed. For the replication cohort (GSE112611), IDAT files were not available from the GEO repository. Instead, the β values file was directly downloaded, along with phenotype data. The downloaded betas used in our analysis were already preprocessed as described in (27).

### Epigenetic Clocks and Other DNAm Estimators of Aging

In this study, we investigated a range of DNAm-based estimators of aging. These included DNAm estimators of biological age, telomere length and the pace of aging. In both the discovery and replication cohorts, values for DNAm GrimAge (7), GrimAge2 (18), PhenoAge (17) and Epigenetic Age (Zhang) (28) were generated by passing DNAm profiles into the Horvath Laboratory’s online DNA Methylation Age Calculator (https://dnamage.genetics.ucla.edu/home). The pace of biological aging was estimated using DunedinPACE (8), following procedures described in Belsky et al. (8). Regarding telomere length estimates, DNAmTL (29) measures were acquired from Horvath’s online calculator while MI-EN TL estimates were generated using the 407-CpG signature and coefficients described previously (30). Additionally, for the paediatric replication cohort, biological age estimates were produced using the supplied CpG signature and coefficient values for the epigenetic clock trained specifically on a paediatric population (31).

Age accelerations are calculated as the residual values of the regression model fitting the epigenetic age estimates on chronological age. The accelerations used for GrimAge, PhenoAge and DNAmTL were available from the Horvath calculator results file i.e., AgeAccelGrim, AgeAccelPheno and DNAmTLAdjAge respectively. Although, the estimates Epigenetic Age (Zhang) and GrimAge2 are available from the Horvath calculator, the accelerations are not - therefore we calculated these directly. Similarly, the residuals were calculated to acquire MI-EN TLAdjAge. As DunedinPACE is a pace of aging over time, we did not residualise this measure. In line with similar studies (32–34), to better compare effect sizes (odds ratios in this case), all DNAm estimator scores were standardised to have a mean of zero and a standard deviation of one before conducting analysis.

### Statistical analysis

We investigated the relationship between each of the 7 DNAm-based estimators, measured at baseline, and a range of binary IBD-related outcomes. This included association analyses for disease course (short recurrence vs. non-recurrence, long recurrence vs non-recurrence) and subtype identification (UC vs. CD, CD vs. Controls and UC vs. Controls). As DNAm can vary with blood cell proportions and potentially confound associations with outcomes, both univariate and multiple regressions were conducted. Multiple regression assessed the relationship between the DNAm estimators and the binary outcome in the presence of the covariates sex, smoking and six estimated measures of blood cell concentrations (monocytes, granulocytes, CD8 T, CD4 T, NK and B cells).

Further to the association analysis, clinicopathological variables at baseline (albumin, C-reactive protein, haemoglobin and disease activity levels) were investigated for their relationship with the DNAm-based estimators. This included correlation analysis and parametric and non-parametric tests of differences. The Mann-Whitney U-test or t-test evaluated differences in continuous variables between binary groups. ANOVA and Kruskal-Wallis tests were utilised to test for differences between continuous input variables and outcome variables with greater than two categories. Follow-up tests in the form of Tukey’s HSD post-hoc test and Dunn’s test were used to assess any significant pairwise differences between categories of the outcome variable for ANOVA and Kruskal-Wallis respectively. Differences in categorical variables were assessed by chi-squared tests. Additionally, ROC curves were generated using CRP/DunedinPACE values and activity status (active/inactive) in each of the UC and CD groups. All analysis was conducted in *Python* (35) using the libraries *statsmodels* (36), scikit-learn (37), *scipy* (38) and *pandas* (39).

### Genome-wide association analysis

Association analyses were conducted using the R statistical programming package version 4.1.2. β values are the ratio of methylated probe intensity and the sum of methylated and unmethylated intensities, ranging from 0 to 1. Differentially methylated positions were obtained using linear regression implemented using the *limma* package (40, 41) i.e., for each CpG site, linear regression was utilized to assess differences in DNAm values between short recurrence and non-recurrence, and long recurrence and non-recurrence. Analyses were conducted while controlling for potential confounders i.e., age, sex, smoking status and estimated blood cell composition (granulocytes, monocytes, CD4 T, CD8 T, natural killer (NK) and B cells). The blood cell compositions were imputed using Horvath Laboratory’s online calculator (https://dnamage.genetics.ucla.edu/home) and the Houseman method (42). Probes were ranked based on p-value with multiple testing accounted for using the Benjamini-Hochberg correction (43).

## Results

### DNA methylation associations with IBD disease course

First, we assessed genome-wide DNAm patterns in IBD disease course within the discovery cohort – specifically short recurrence vs. non-recurring and long recurrence vs. non-recurring using a methylome-wide association study. One CpG site (cg03583111) was significantly differentially methylated after correction for multiple testing (Benjamini Hochberg (BH) adjusted p-value < 0.05) in IBD patients with long recurrence compared to non-recurrence. No CpG site was significantly differentially methylated after correction for multiple testing in IBD patients with short recurrence compared to non-recurrence. Tables S1 and S2 in the supplementary information (SI) show the top 10 differentially methylated positions (DMP) for long and short recurrence respectively.

### IBD patients exhibit epigenetic age acceleration compared to controls

Given previous reports of epigenetic age acceleration in IBD patients compared with controls (15, 16), we examined this relationship in the discovery cohort. DunedinPACE, GrimAge and GrimAge2 showed significant associations with IBD/control status (p=0.003, p=0.004 and p=0.008 respectively). Associations remained significant after correction for multiple testing (BH). After covariate adjustment, four of the seven investigated DNAm aging signatures showed evidence of association but did not remain significant after multiple testing correction (Table S3). The mean DNA methylation levels for each of the three significant DNAm aging signatures (Figure 1) indicate that individuals with IBD exhibited significantly increased biological aging compared with controls. Figure S1 shows a plot of the odds ratios corresponding to each DNAm signature of aging in relation to IBD/control status.

**Figure 1:**
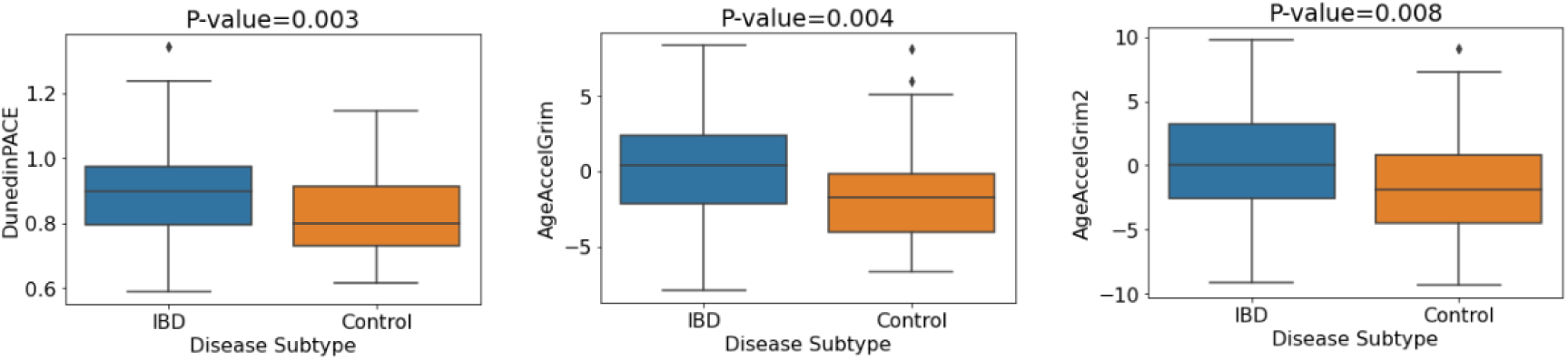
Mean values of DNA methylation aging signatures for IBD vs. control status. P-values pertain to the logistic regression model without covariate adjustment.

Furthermore, significant differences in first- and second-generation epigenetic clocks were reported between CD patients and non-IBD controls (15, 16). Therefore, we examined this relationship in our discovery cohort. DunedinPACE, GrimAge and GrimAge2 showed significant associations with CD/control status (p=0.0006, p=0.0006 and p=0.0019 respectively). Associations remained significant after correction for multiple testing (BH) and adjustment for covariates (Table S4). In addition, both DNAm-based telomere length estimators investigated, DNAmTL and MI-EN TL, showed evidence of association with CD. Table S4 contains the p-values and odds ratios (plotted in Figure S2) for each of the DNAm signatures of aging relating to CD vs. control status associations in the discovery cohort. Individuals with CD exhibit significantly increased biological aging compared with controls (Figure 2).

**Figure 2:**
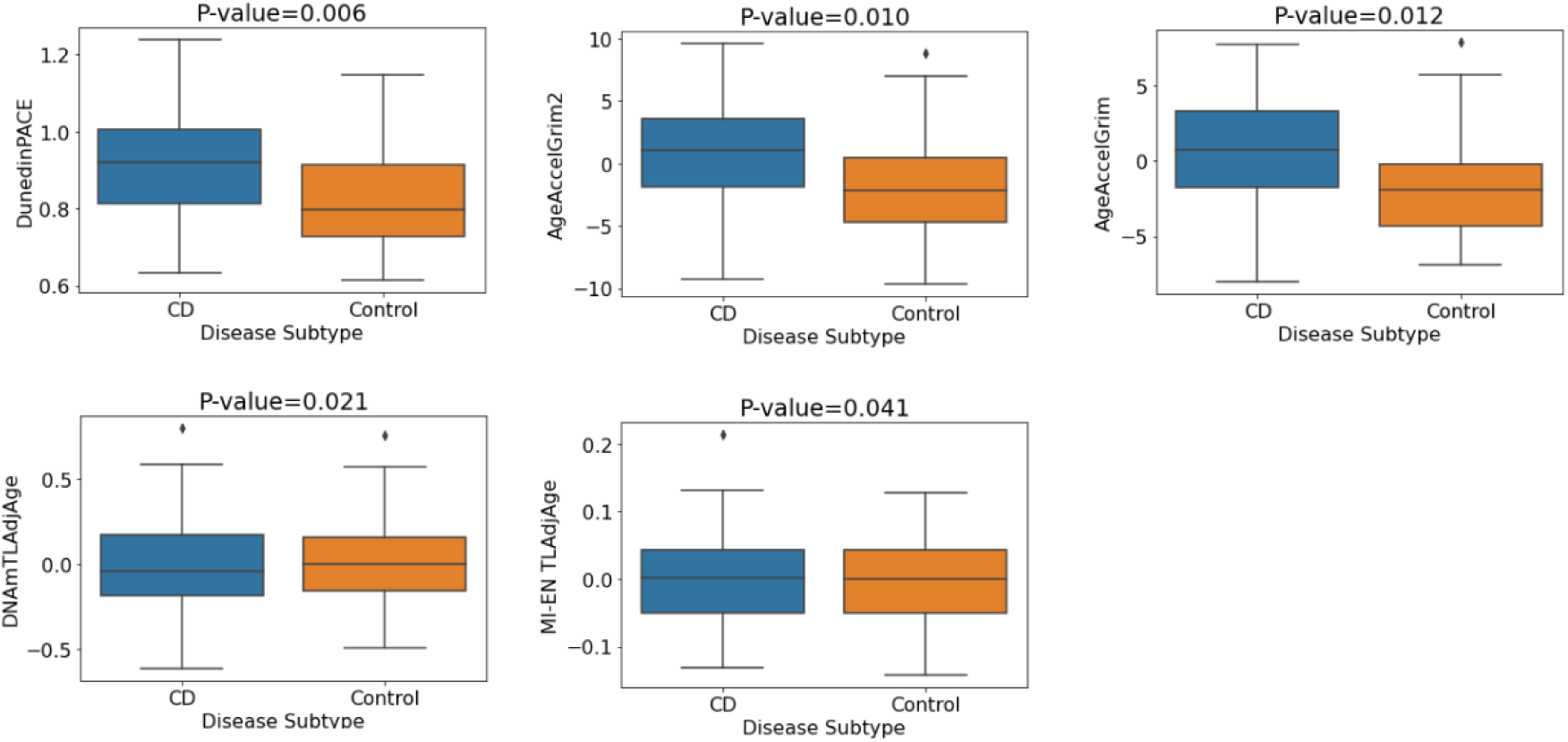
Mean values of DNA methylation aging signatures for CD vs. control status. P-values pertain to the multiple logistic regression model with covariate adjustment (sex, smoking, blood cell concentrations).

We also investigated associations of DNAm estimators of aging in UC patients vs. controls. One estimator, DNAmTL, showed evidence of association (p=0.038) but did not remain significant after multiple testing correction. No other significant associations were observed (Table S5).

### Replication of IBD-associated epigenetic age acceleration in independent cohorts

Next, we aimed to replicate our association between the DNAm signatures of aging and IBD using two independent publicly available IBD cohorts (GSE87648 (23) and GSE112611 (27)). Given the consistency of significant associations apparent in the discovery cohort for DunedinPACE, GrimAge and GrimAge2, we focused replication of results on these three epigenetic clocks. The first replication cohort (GSE87648) included 175 controls and 204 IBD cases. Investigating the relationships between IBD and control status, we observed significant associations between the three clocks and IBD/control status (DunedinPACE: p-value=8.08 x 10^-21^, GrimAge2: p-value=1.11 x 10^-15^, GrimAge: p-value=2.72 x 10^-8^). P-values remained highly significant after multiple testing correction (BH) and after adjusting for covariates. Again, we observed increased epigenetic age in IBD patients compared with controls (Figure 3). Table S6 contains the p-values and odds ratios (plotted in Figure S3) relating to IBD/control status associations for both adjusted and unadjusted models.

**Figure 3:**
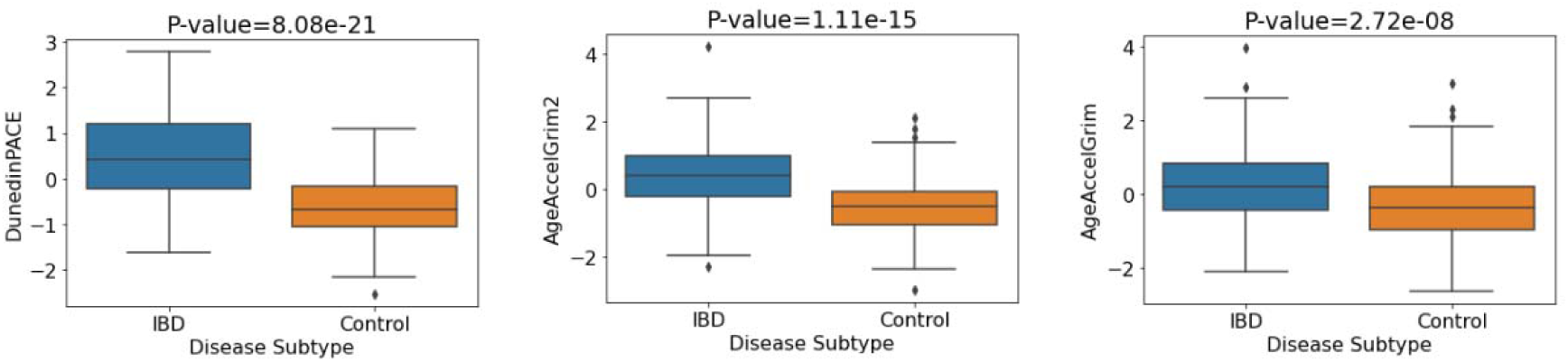
Mean DNA methylation values of aging signatures for IBD vs. control status. P-values correspond to the unadjusted logistic regression model.

Next, we investigated the relationships between the three epigenetic clocks and both CD/control status and UC/control status. Firstly, comparing CD and controls cases, the analysis involved 278 participants comprising 175 controls and 103 CD cases. Significant associations were observed between each of DunedinPACE, GrimAge and GrimAge2 with CD/control status (DunedinPACE: p-value=7.01 x 10^-17^, GrimAge2: p-value=2.23 x 10^-14^, GrimAge: p-value=6.65 x 10^-9^). All p-values remained significant after BH correction. Strongly significant associations persisted on controlling for covariates (sex and blood cell concentrations), with significantly increased biological aging evident in CD patients compared to controls (Figure 4). The relevant p-values and odds ratios for each of the DNAm signatures of aging are shown in Table S7.

**Figure 4:**
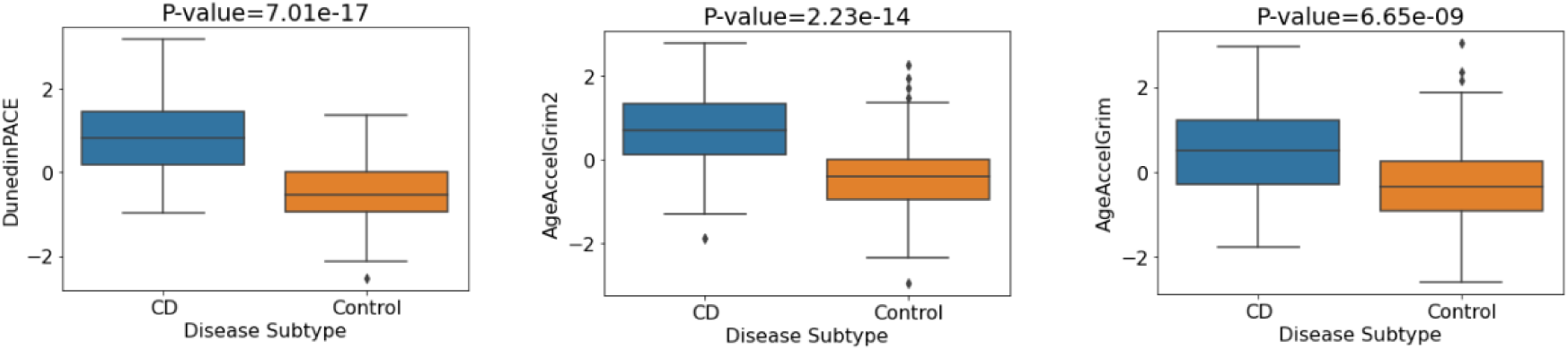
Mean values of DNA methylation aging signatures for CD vs. control. P-values are for the logistic regression model without covariate adjustment.

Next, we compared UC cases (N=101) and controls (N=175) for the three epigenetic clock measures. We observed significant associations for each of DunedinPACE (5.75 x 10^-14^), GrimAge2 (8.91 x 10^-9^) and GrimAge (5.54 x 10^-4^) with UC/control status. Associations remained significant after multiple testing correction and after adjusting for potential confounders (sex, blood cell concentrations). The mean methylation levels of each epigenetic clock (Figure 5) indicate higher epigenetic age in UC patients versus controls. Table S8 contains the p-values and odds ratios (plotted in Figure S5) relating to UC/control status associations for both adjusted and unadjusted models.

**Figure 5:**
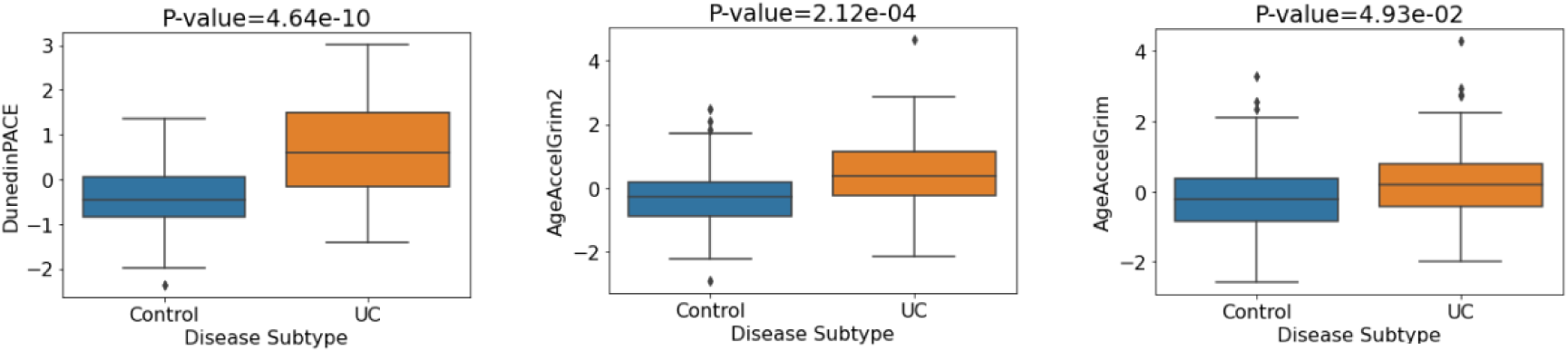
Mean values of DNA methylation aging signatures for UC vs. control. P-values correspond to the adjusted logistic regression model (sex, blood cell concentrations).

A second replication cohort was tested for CD-associated changes in epigenetic clock acceleration. The paediatric CD only cohort (GSE112611) was composed of 238 participants (164 CD cases and 74 controls). As existing epigenetic clocks are predominantly adult-oriented, we applied a DNAm-based biological age estimator for children (31) in addition to both GrimAge clocks and DunedinPACE. For the regression model with no covariates, we observed significant associations between each of the four tested clocks and CD/control status (AgeAccelGrim2: p-value= 1.34e^-14^, DunedinPACE: p-value= 2.04e^-14^, GrimAge: p-value=6.67e^-14^, Wu Age (Paediatric) AdjAge: p-value= 1.59e^-02^). All associations were significant after multiple testing correction. Regarding effect size, each standard deviation increment in AgeAccelGrim2 (OR=8.63, CI 4.99, 14.94) and DunedinPACE (OR=22.46, CI 10.12, 49.87) was related to an 8-fold and 22-fold increase in likelihood of being a CD subject as opposed to a control subject respectively (Figure S6). Highly significant associations were also observed when controlling for covariates i.e., sex and blood cell concentrations. P-values and odds ratios for each of the DNAm signatures of aging are shown in Table 6 with mean DNA methylation levels for each signature (Figure 6) indicating CD patients’ increased biological aging versus controls.

**Figure 6:**
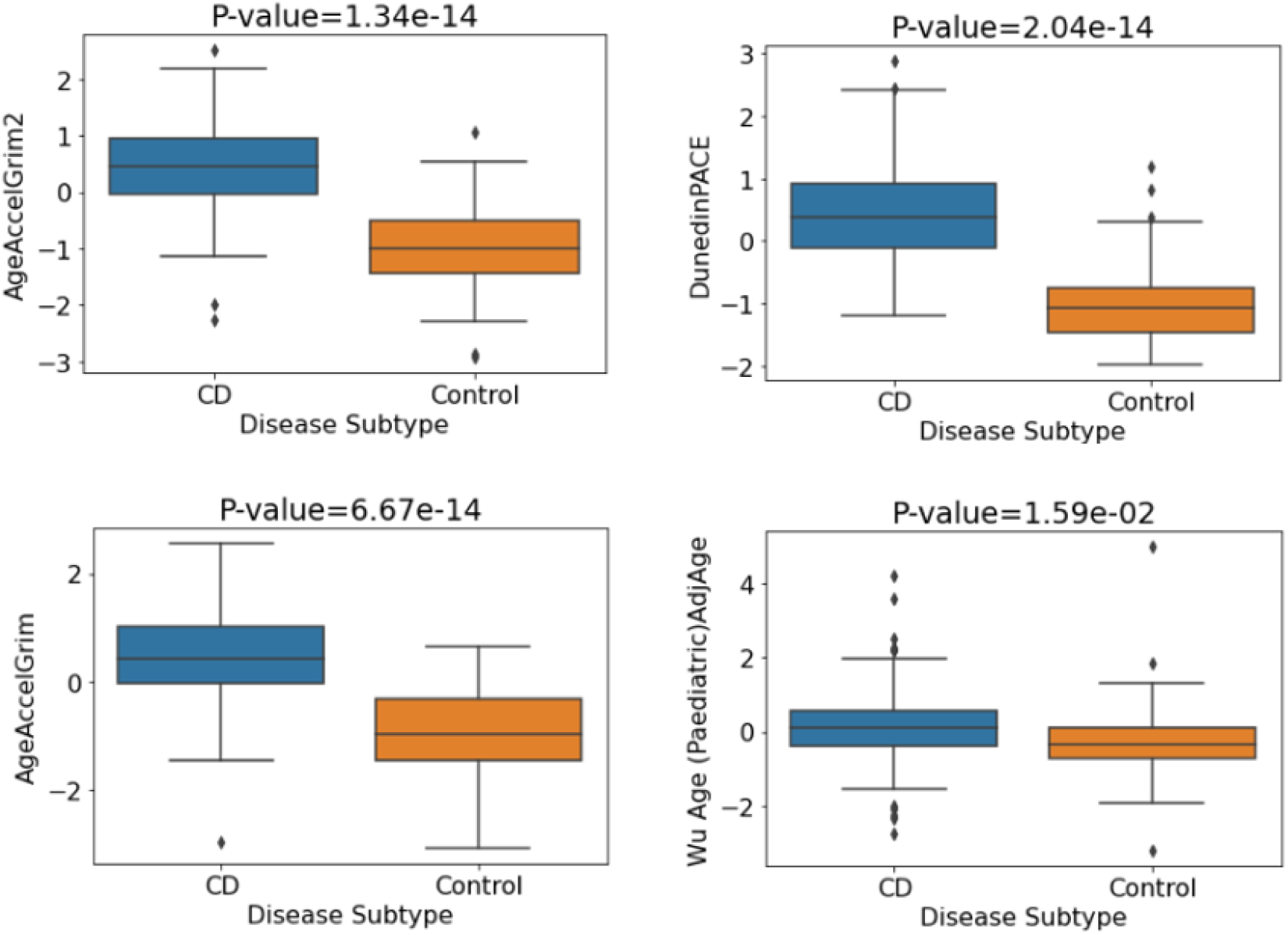
Mean values of DNA methylation aging signatures for CD vs. control. P-values are for the logistic regression model with no covariate adjustment.

**Table 6:**
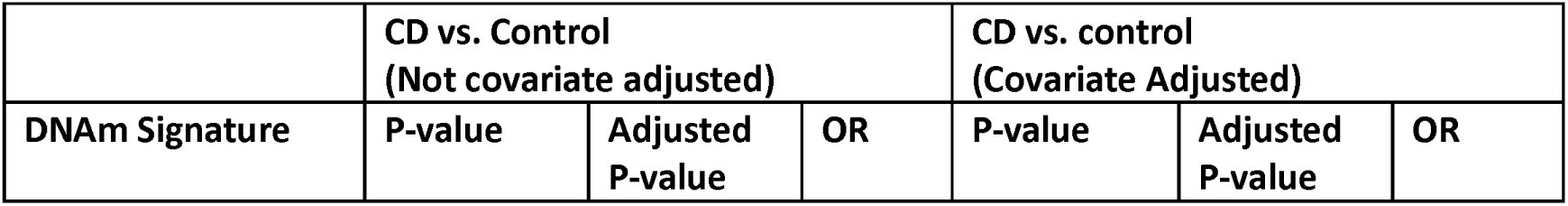
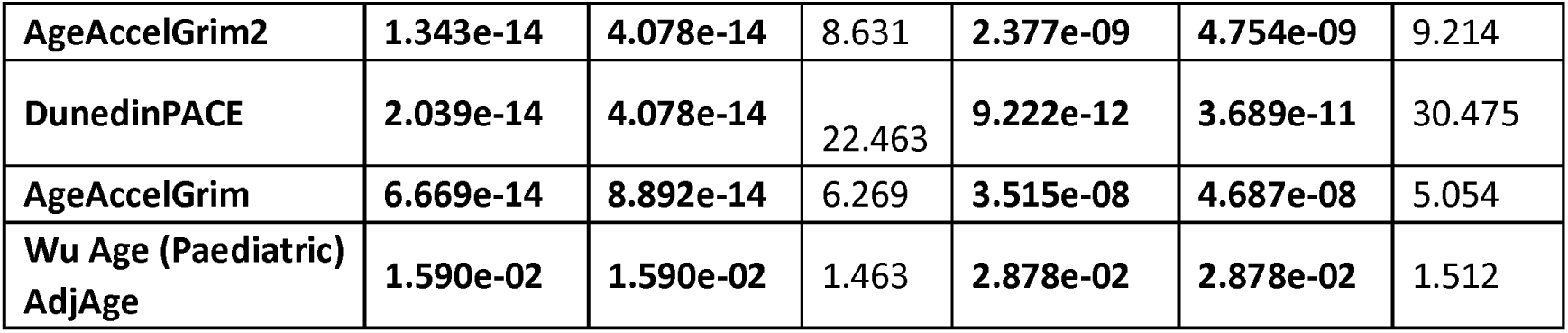
Association of standardised DNA methylation aging signatures with CD/control status for both unadjusted and adjusted logistic regression models (sex, blood cell concentrations) for the GSE112611 replication data set.

| DNAm Signature | CD vs. Control<br>(Not covariate adjusted) |  |  | CD vs. control<br>(Covariate Adjusted) |  |  |
| --- | --- | --- | --- | --- | --- | --- |
|  | P-value | Adjusted P-value | OR | P-value | Adjusted P-value | OR |
| AgeAccelGrim2 | 1.343e-14 | 4.078e-14 | 8.631 | 2.377e-09 | 4.754e-09 | 9.214 |
| DunedinPACE | 2.039e-14 | 4.078e-14 | 22.463 | 9.222e-12 | 3.689e-11 | 30.475 |
| AgeAccelGrim | 6.669e-14 | 8.892e-14 | 6.269 | 3.515e-08 | 4.687e-08 | 5.054 |
| Wu Age (Paediatric)<br>AdjAge | 1.590e-02 | 1.590e-02 | 1.463 | 2.878e-02 | 2.878e-02 | 1.512 |

### DNA methylation signatures of aging and their association with disease course in IBD

We examined the relationship between DNAm signatures of aging and disease course in IBD patients in the discovery cohort using logistic regression. We observed evidence of association between one epigenetic clock, DunedinPACE, and short recurrence status (p=0.032) – individuals whose IBD recurred in the short-term (1 year) exhibited faster biological aging (DunedinPACE scores), at baseline, than those who did not clinically relapse (i.e. no clinical recurrence of disease activity requiring treatment escalation over short (treatment escalation in the first study year) and long term (after the first year of study). However, this association did not remain significant after stringent correction for multiple testing. Regarding long-term recurrence, no DNAm signature of aging exhibited a significant difference between those who experienced clinical relapse and those who did not, over the assessed 8-year period. Table S9 outlines the p-values and odds ratios for each of the DNAm signatures of aging for the short and long-term recurrence associations (not adjusted for covariates).

Next, we investigated associations using a multiple regression model adjusted with the covariates sex, smoking and a range of white blood cell concentrations to control for potential confounding (28). Once again, only DunedinPACE (p=0.036) exhibited evidence of association with short-term IBD recurrence in the multiple regression model adjusted for covariates. Moreover, each unit increase (standard deviation) in standardised DunedinPACE (OR=1.73, CI 1.04, 2.89) was related to more than a 70% greater likelihood of IBD short-term recurrence (Figure 7), however this association did not remain significant after stringent adjustment for multiple testing. No evidence of association was observed for long recurrence. Table S10 shows the p-values and odds ratios pertaining to each of the DNAm aging signatures for the short and long-term recurrence associations (adjusted for covariates).

**Figure 7:**
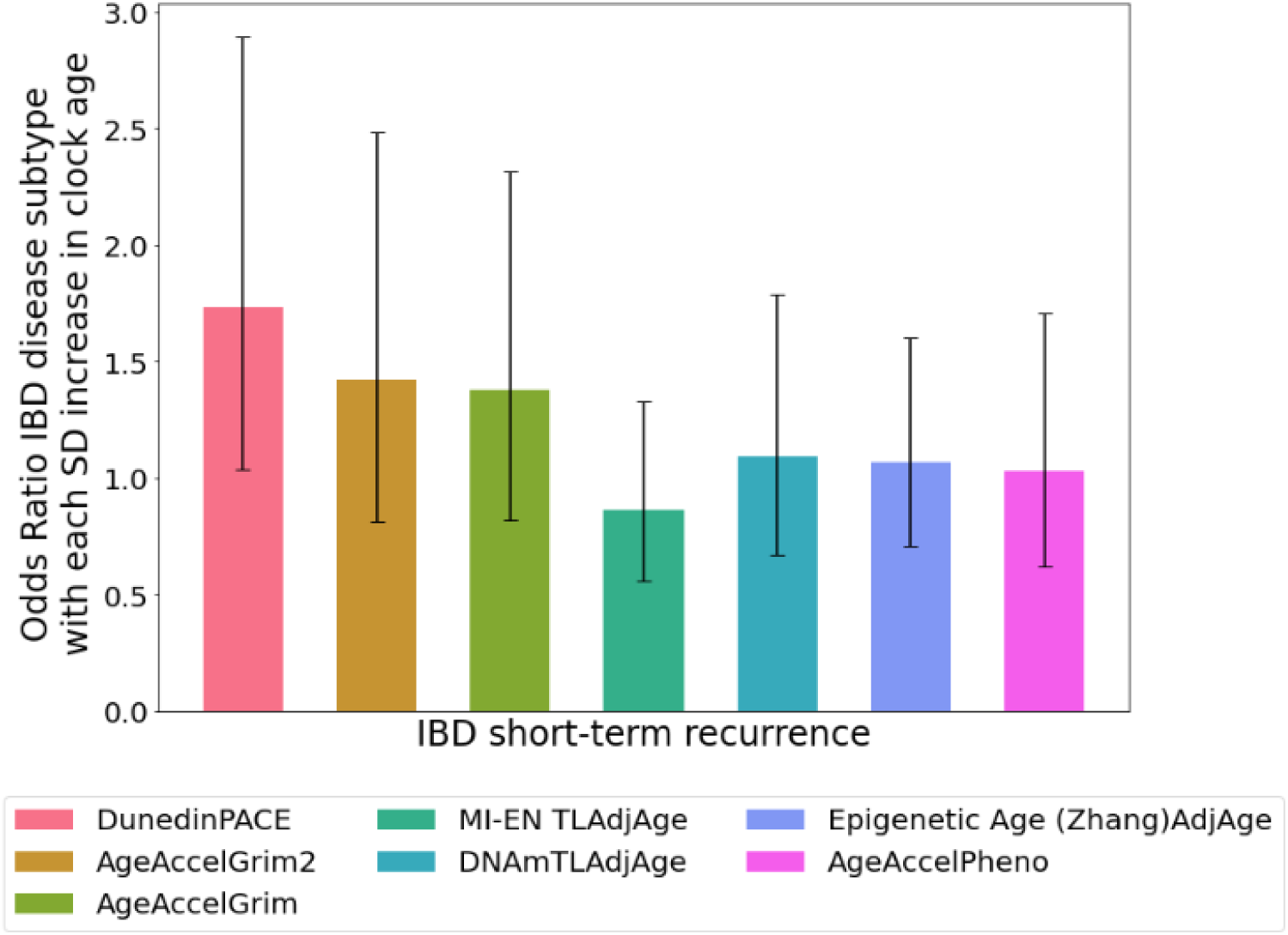
Odd ratios of DNAm signatures of aging in blood for IBD disease course (short-term recurrence). Odds ratios adjusted for sex, smoking status and white blood cell concentrations (n=146). Error bars indicate 95% confidence intervals.

### DNA methylation signatures of aging are more strongly associated with CD than UC

To explore the relationship between epigenetic age acceleration and IBD subtypes, we analysed a range of DNA methylation clocks and found significant associations between the clocks GrimAge, GrimAge2, DunedinPACE and PhenoAge with disease subtype (p=0.018, p=0.036, p=0.037 and p=0.040 respectively). After adjustment for covariates, a further signature, MI-EN TLAdjAge (p-value=0.032), became significant. The five signatures remained significant after adjustment for covariates and BH correction (see Table 7). The boxplots in Figure 8 indicate the mean values of the DNAm aging signatures across disease subtype. Considering the plotted mean values, individuals with CD exhibited significantly higher biological aging or significantly shorter estimated telomere length (MI-EN TLAdjAge) compared with UC patients. P-values and odds ratios for associations of each of the DNAm signatures of aging with disease subtype are shown in Table 7 while the odds ratios are plotted for each estimator in Figure S7. We then replicated these findings in an independent cohort (GSE87648, N=204 (101 UC and 103 CD cases)). Significant associations were found for GrimAge clocks (GrimAge2: p=0.008, GrimAge: p=0.015), which remained significant after adjustment and BH correction (Table 7). CD subjects consistently showed increased biological aging compared to UC subjects (Figure 9).

**Table 7:**
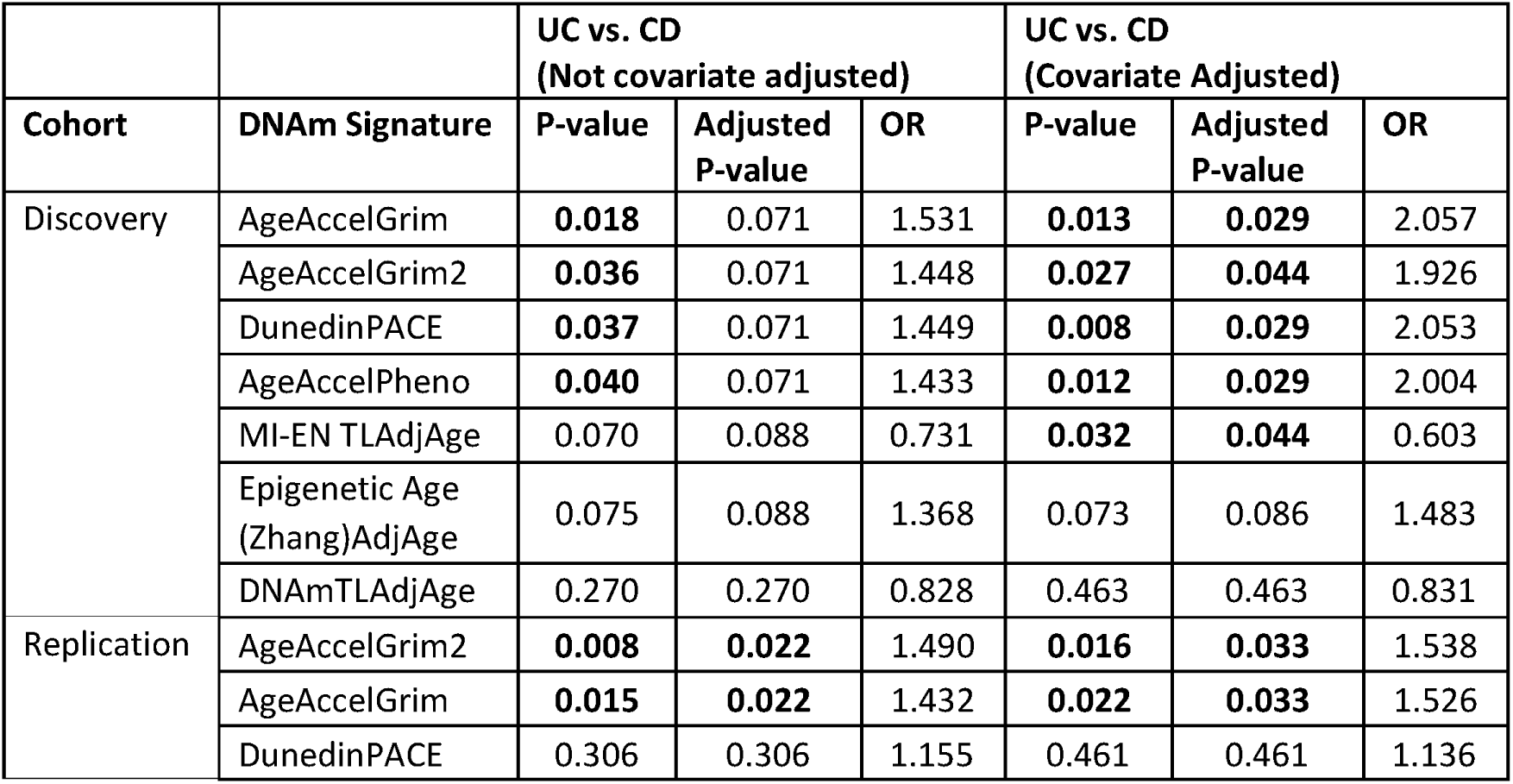
Association of standardised DNAm aging signatures with disease subtype (UC vs. CD) for the unadjusted and adjusted (sex, smoking, blood cell concentrations) logistic regression models – discovery and replication (GSE87648) cohorts.

**Figure 8:**
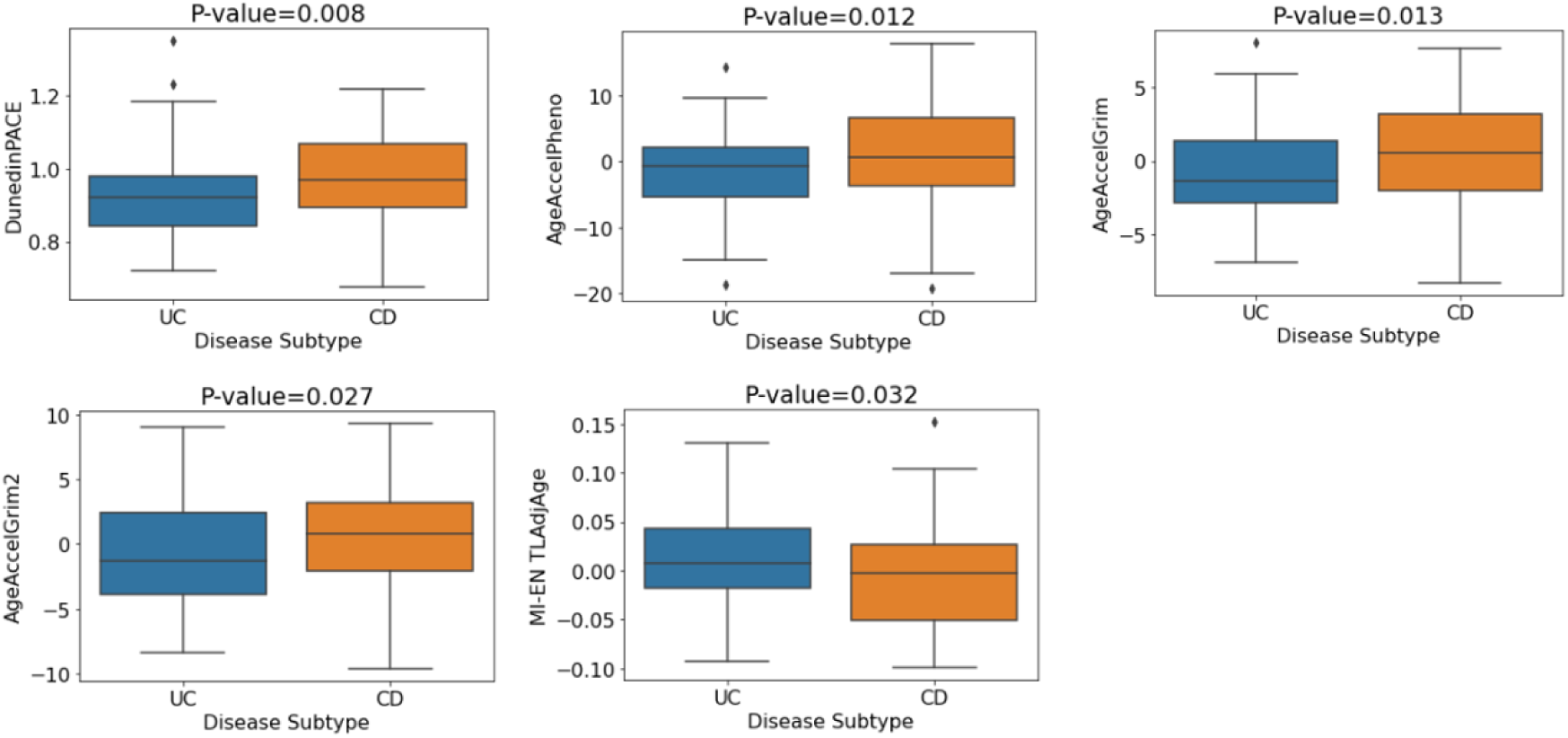
Mean DNA methylation values of aging signatures for disease subtype (discovery cohort). P-values are for the multiple logistic regression model with covariate adjustment (sex, smoking, blood cell concentrations).

**Figure 9:**
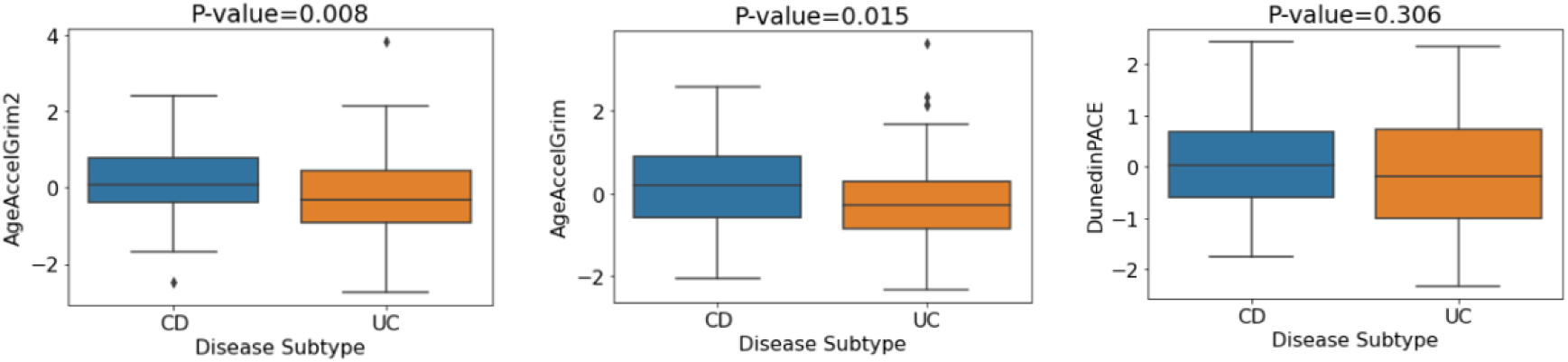
Mean values of DNA methylation aging signatures for disease subtype (GSE87648 replication cohort). P-values pertain to the logistic regression model without covariate adjustment.

### DunedinPACE is associated with inflammatory indices and disease activity in UC patients

In our discovery cohort, clinical measures of inflammatory indices (C-reactive protein (CRP), Albumin and Haemoglobin) and disease activity measures at baseline were available. We found moderate correlations in our discovery cohort between inflammatory indices and a range of epigenetic clocks. Figure 10 outlines the results of a correlation analysis between C-reactive protein (CRP), albumin and haemoglobin, and a range of DNAm-based estimators for both UC and CD patients. DunedinPACE exhibits moderate correlations with CRP (r=0.45, p=4.4e-04) and albumin (r=-0.54, p=1.3e-05) in UC patients and weak correlations with CRP (r=0.23, p=0.044) and albumin (r=-0.27, p=0.016) in CD patients. Additionally, AgeAccelGrim is weakly correlated with both CRP (r=0.21) and albumin (r=-0.28) in UC patients.

**Figure 10:**
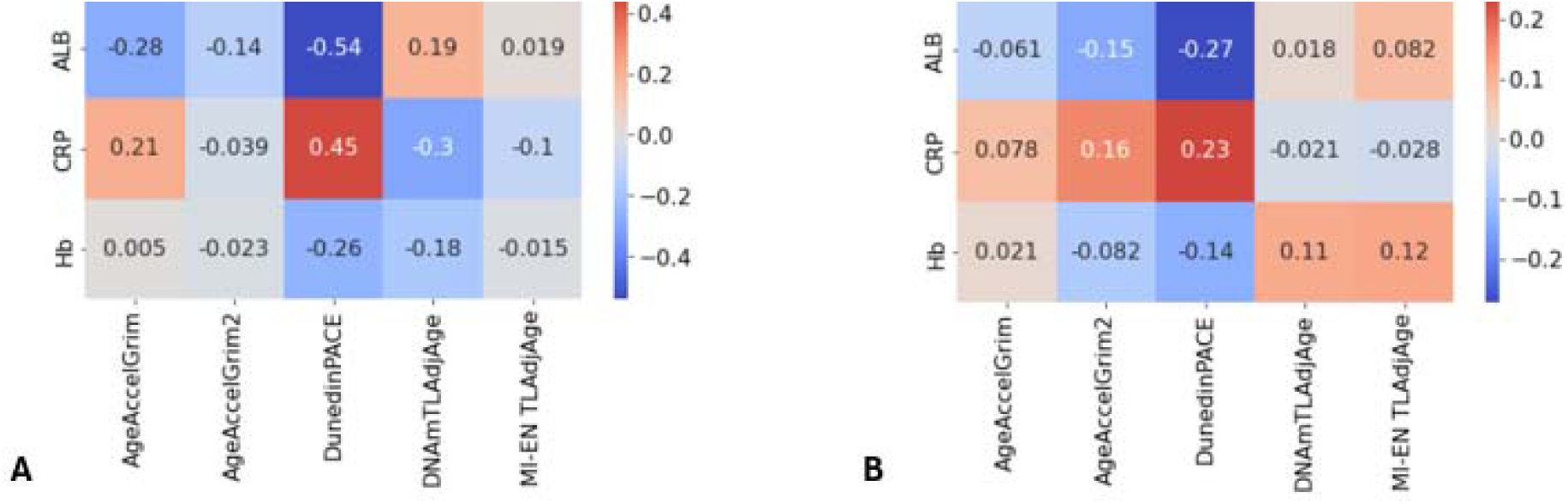
Correlation heatmaps of clinicopathological variables of IBD patients and DNAm-based signatures. (**A**) UC and (**B**) CD patients in the discovery cohort. CRP: C-reactive protein, ALB: albumin, Hb: haemoglobin.

In our discovery cohort, disease activity measures at baseline were available for both UC and CD in the form of both a binary variable (Active/Inactive) and a more granular 4-category variable (Inactive/Mild/Moderate/Severe). Firstly, we examined associations between DNAm signatures of aging and binary disease activity in IBD patients (N=146). As such, significant group differences were observed in both DunedinPACE and AgeAccelGrim2 - with the active disease group associated with a higher pace of aging (DunedinPACE (t=3.169, p=0.002)) and higher age acceleration (AgeAccelGrim2 (t=1.977, p=0.05)) compared to the inactive group. Next, we examined associations in both IBD subtypes. For UC, significant group differences existed in both DunedinPACE and AgeAccelGrim. The active disease group was associated with higher age acceleration (GrimAge (U=669, p=0.003)) and higher pace of aging (DunedinPACE (t=3.233, 0.002)) compared to the inactive group. Finally, in CD patients no significant differences were observed for any of the DNAm aging signatures and disease activity. For UC patients, Figures 11 and 12 show the DNAm aging signature levels for the binary and four-category activity states.

**Figure 11:**
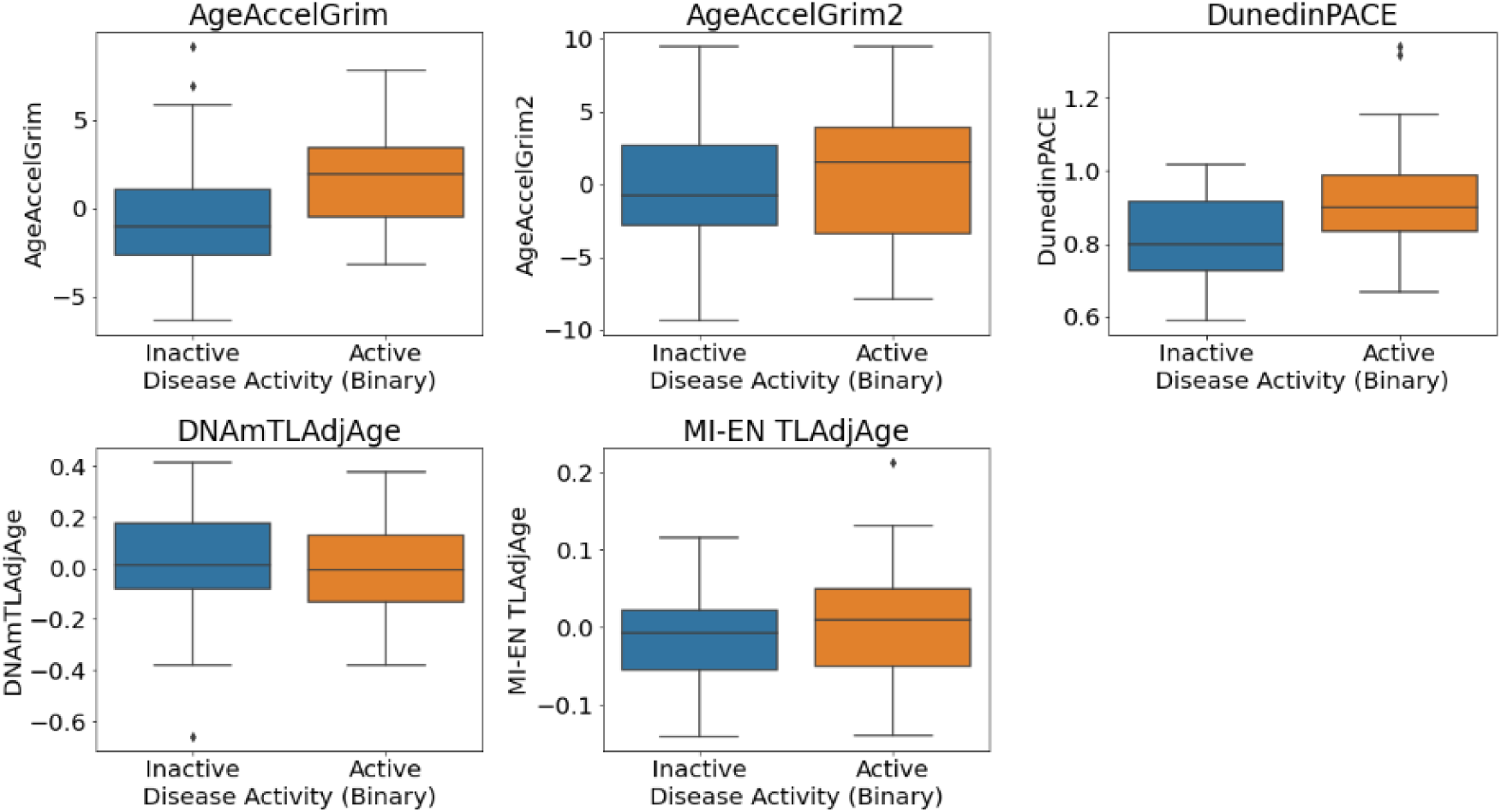
Disease activity (Active/Inactive) in UC patients in the discovery dataset (n=61).

**Figure 12:**
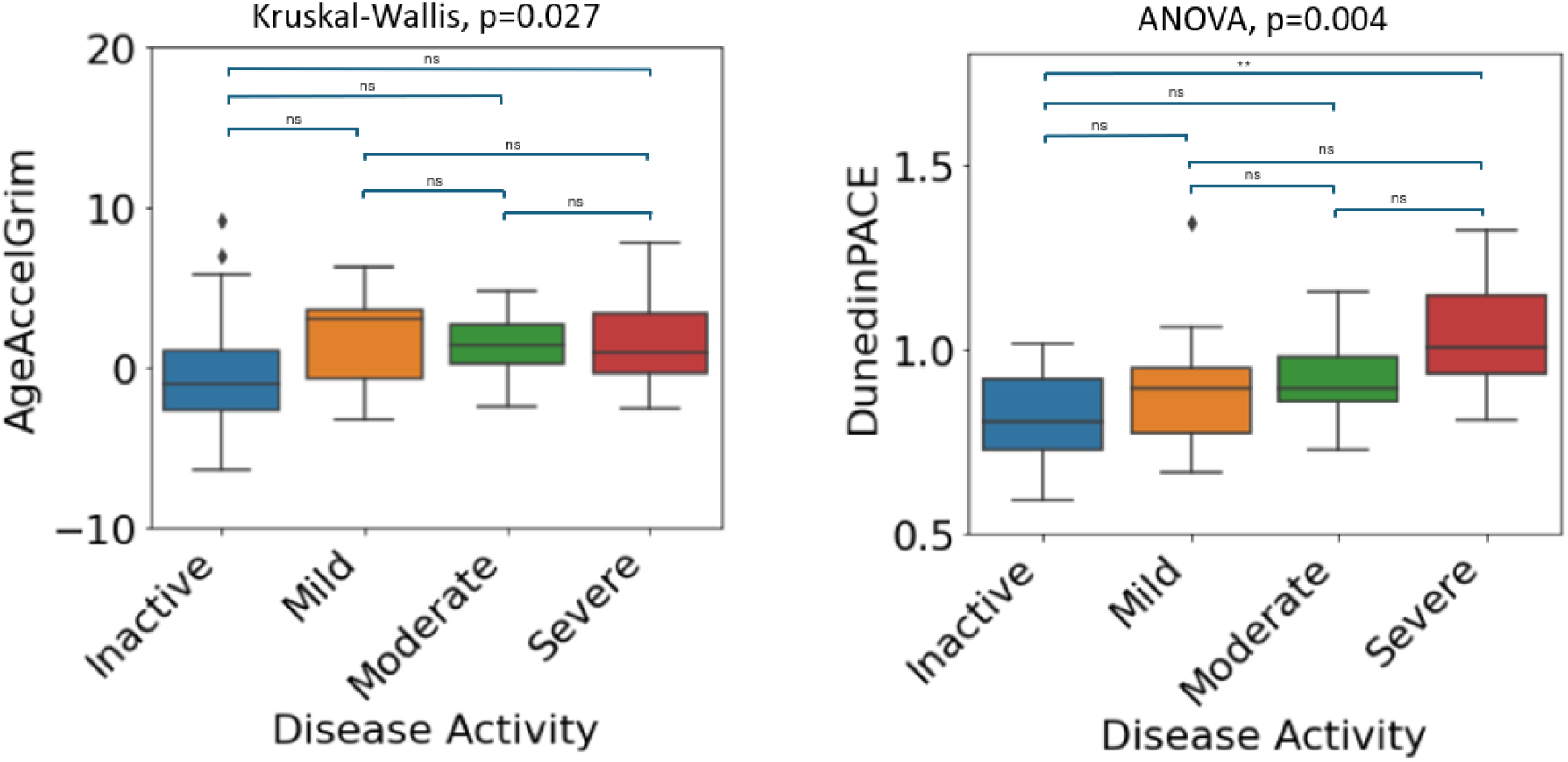
Four-category disease activity for AgeAccelGrim and DunedinPACE in UC patients in the discovery dataset (n=61). Significance codes: *: <0.05, **: <0.01, ***: <0.001, ns: not significant at 0.05.

Given the observed significant relationship between both AgeAccelGrim and DunedinPACE with binary disease activity in UC patients, we further investigated both epigenetic clocks for the 4-category disease activity variable. Levels of AgeAccelGrim (Kruskal-Wallis p=0.027) and DunedinPACE (F(3,57)=5.044, p=0.004) tend to increment across groups with increasing severity of disease activity (Figure 12).

### Comparison of DunedinPACE and CRP as biomarkers for disease activity

To assess DunedinPACE as a potential biomarker of disease activity in IBD, we evaluated the efficacy of both CRP and DunedinPACE in discriminating disease activity in the discovery cohort using an ROC analysis. Considering CRP and DunedinPACE for IBD patients – both measures have similar areas under the ROC curve (AUC) i.e., AUC=0.68 and 0.66 respectively. We further investigated DunedinPACE in both the UC and CD patient subsets. In CD, DunedinPACE did not perform as well as CRP in discriminating active and inactive disease (AUC=0.6 and 0.76 for DunedinPACE and CRP respectively (Figure 13)). However, in UC, Figure 13 indicates the superior performance of DunedinPACE over CRP in classifying disease activity. DunedinPACE achieves an AUC of 0.71 in comparison to CRP’s AUC of 0.57. In the case of UC, a DunedinPACE cut-off value of 0.93 corresponds to a sensitivity of 69.5% and specificity of 68.7%.

**Figure 13:**
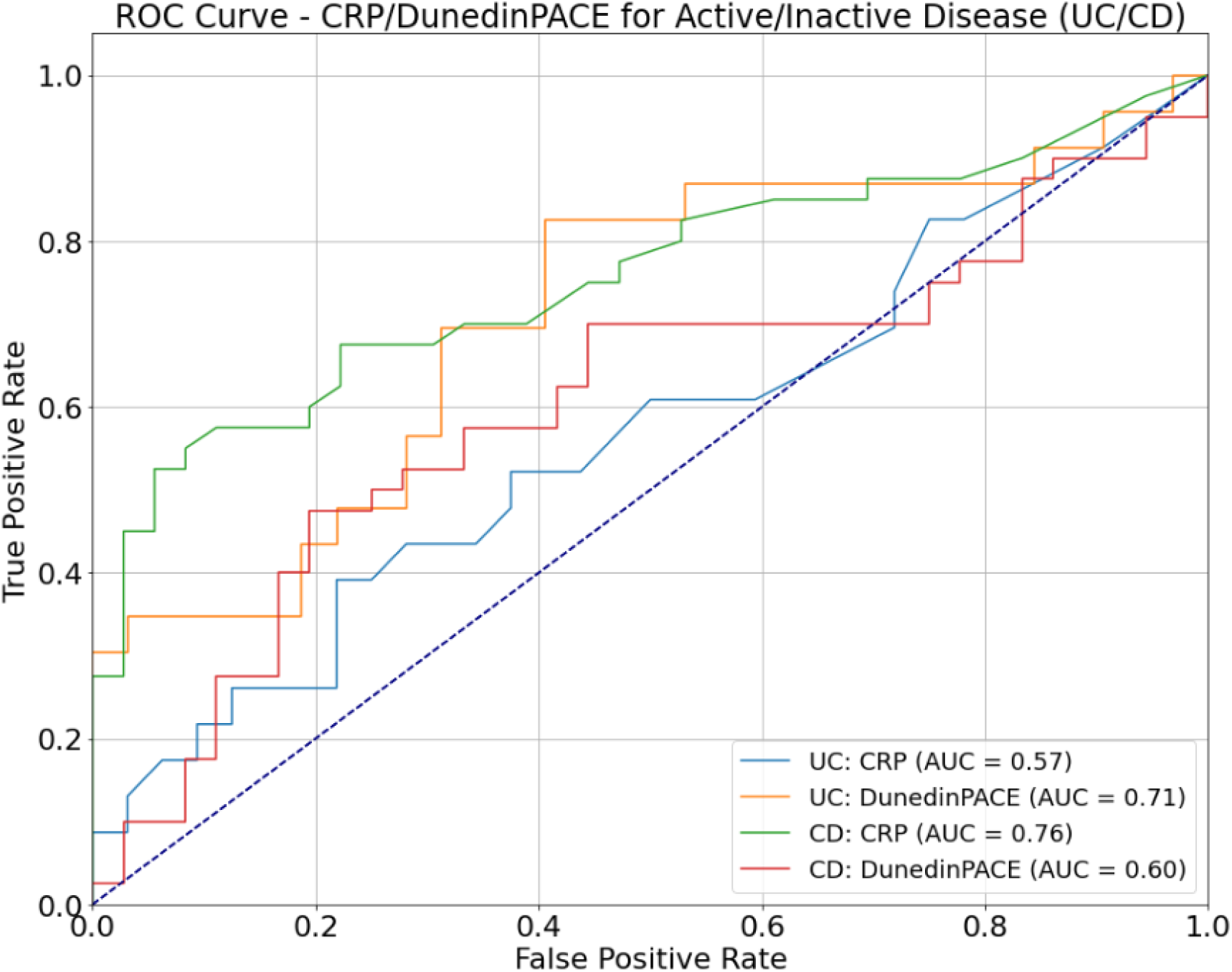
ROC curve analysis of DunedinPACE versus CRP. ROC curves show performance of CRP and DunedinPACE measures in discriminating binary disease activity (Active/Inactive) for UC and CD patients in the discovery cohort.

## Discussion

IBD is a debilitating proinflammatory condition that often results in the development of age-associated comorbidities (including physical frailty), leading to reduced quality of life and ultimately increased mortality (44). Aging is also associated with a chronic increase in circulating proinflammatory cytokines and a decrease in the level of anti-inflammatory cytokines, a process referred to as ‘inflammageing’ (45). Patients with IBD have a shorter life expectancy and an increased risk of developing aging-related diseases such as cardiovascular disease, dementia, and type 2 diabetes (46). To date numerous studies have investigated epigenetic clock age in health and disease and have shown marked differences in disease and mortality among individuals of equivalent chronological age (32, 47–51). Recent studies have identified a significant correlation between epigenetic age acceleration (defined as the discrepancy between predicted age, as determined by DNA methylation patterns, and chronological age) and IBD, as well as its subtypes (15, 16).

The present study investigated whether the rate of biological aging in blood is increased in IBD patients compared to controls. We report strong evidence for accelerated DNA methylation pace of aging (DunedinPACE), in IBD patients compared to controls, replicating our findings in an independent adult IBD and paediatric IBD cohort. Moreover, the DunedinPACE epigenetic clock is the only clock that is positively associated with short-term IBD recurrence (within 1 year) compared to those without clinical relapse (no need for treatment escalation in both the short and long term (up to 8 years)). DunedinPACE, a novel blood biomarker that reflects aging pace over two decades, differs from other epigenetic clocks by capturing the rate of system deterioration. It is linked to increased risks of morbidity, disability, and mortality, and has shown faster aging in young adults exposed to childhood adversity (8). Future research, in a longitudinal study of IBD patients, is required to evaluate the potential of DunedinPACE as a novel blood-based biomarker of disease recurrence in IBD patients.

Next, we examined the association of the epigenetic clocks with a range of clinicopathological variables, including measures of inflammation (CRP and albumin) and disease activity. Disease activity, measured using a combination of clinical, biological and endoscopic variables, was available for both UC and CD in addition to measures of disease severity (Mild/Moderate/Severe) in the discovery cohort. Firstly, we assessed associations between DNAm signatures of aging and disease activity in IBD patients where, in comparison to the inactive group, the active disease group was associated with higher age acceleration (AgeAccelGrim2 (p=0.05)) and a higher pace of aging (DunedinPACE (p=0.002)). Next, we examined the IBD subtypes (UC and CD) separately to establish if there were any differences in the associations between the epigenetic clocks and disease activity. In UC patients, both DunedinPACE and AgeAccelGrim were significantly associated with disease activity. The active disease group was associated with higher age acceleration (GrimAge (U=669, p=0.003)) and higher pace of aging (DunedinPACE (t=3.233, 0.002)) compared to the inactive group. Moreover, levels of AgeAccelGrim and DunedinPACE tend to increment across groups with increasing severity of disease activity. In contrast, in the CD patient group, no significant differences were observed for any of the epigenetic clocks and disease activity in CD patients. Taken together, these results suggest that the epigenetic age acceleration differences that we observed in CD patients compared to UC patients are not driven by variation in disease activity between the groups.

The above findings suggest that alternative disease pathways might be influenced in the two IBD subtypes. Additionally, the finding that patients with active UC exhibit increased epigenetic age acceleration similar to CD patients (irrespective of disease activity status) compared to inactive UC patients, is in contrast to previous studies by our group and others, which show no site-specific differences in whole-blood DNAm profiles of UC and CD patients with active disease (3, 16). This highlights the power of the next generation of epigenetic clocks, which have been trained and tested on large datasets, and which are composite multisystem biomarkers, to act as proxies for system integrity (52).

In addition, to examining associations with disease activity, we also investigated the relationship between the epigenetic clocks and measures of inflammation (CRP and Albumin) in our discovery cohort. Serum CRP is one of the best-studied non-invasive biomarkers of inflammation in IBD (53) while serum albumin is a negative acute-phase reactant widely applied to assess the inflammatory condition of the human body (54). Of interest, DunedinPACE exhibited the strongest correlation with CRP and albumin in UC patients, with weaker associations observed in CD patients. In contrast, previous studies by Kalla et al. (16) and Ventham et al. (15) found poor correlation between epigenetic age acceleration and inflammatory indices in IBD. This may partly reflect the biomarkers underlying DunedinPACE which include CRP. Additionally, DunedinPACE is distinct from other DNAm clocks, which estimate the progress of aging leading up to a cross-sectional point in time, as it instead reflects a rate of decline (8).

In addition, we wanted to compare the predictive ability of our strongest performing epigenetic clock (DunedinPACE) with the IBD biomarker, CRP. CRP has a short half-life, making it a valuable marker for the detection of disease activity in IBD. In contrast to CD, UC has a modest to null CRP response despite active inflammation (55). However, several studies have shown CRP’s ability to discern active from inactive disease in UC – reporting AUCs, sensitivities and specificities in the ranges 0.6 - 0.81, 62.2% - 69.4% and 53% - 97% respectively (56–58). In contrast, DunedinPACE estimates of the pace of aging discriminate activity in UC patients with an AUC, sensitivity and specificity of 0.71, 69.5% and 68.7% respectively, highlighting its potential as a useful biomarker of activity in UC.

### Limitations and Challenges

Our study does have several caveats, including the relatively small (by epidemiological standards) sample size. However, we replicated our IBD and subtype epigenetic age accelerations associations in two independent replication cohorts. Furthermore, PBMC samples are heterogeneous, composed of multiple cell types with distinct DNAm profiles. This can lead to potential confounding of results given that cell type proportions can vary from sample to sample (59). In this study, we used algorithms that estimate cell counts, which are commonly used in studies to adjust for differential cell proportions by adding these estimates as covariates to statistical models. A further caveat is that our study group was heterogeneous with regard to disease location, biologic use and disease duration. However, this heterogeneous cohort allowed us to determine epigenetic similarities and differences between those with CD and UC, and also compare DNAm-based estimators of patients with active and inactive IBD. However, larger, more powerful studies on more homogeneous patient populations with respect to disease duration, disease location and other clinical features will be required to tease out these associations more thoroughly. Additionally, it will be important to further consider the impact of medications used to treat IBD on biological aging.

Our findings suggest that current active disease is not a driving force in epigenetic age acceleration or pace of aging observed in CD patients, thus additional research is required to tease apart the molecular underpinnings of this association. Although inflammation subsides after successful treatment of IBD, substantial acute intestinal inflammation persists in the majority of patients with clinically quiescent CD (60). Furthermore, it has previously been indicated that clinical disease activity is associated with mucosal inflammation in UC patients but not in CD patients (61). Given that CRP is an underlying factor in DNAm aging signatures such as GrimAge2 and DunedinPACE, these biomarkers partially account for inflammatory states in individuals. This supports our findings wherein we observed significant differences in epigenetic age and pace of aging for active and inactive UC patients but not for CD patients. Previous research suggests that CD is marked by systemic inflammation, whereas UC as fluctuating inflammatory responses. Taken together, these findings suggest that blood-based DNAm signatures could serve as biomarkers for detection and monitoring in IBD patients.

## Conclusions

This study provides important new evidence that DunedinPACE, a biomarker for the pace of aging based on organ-system integrity, may have utility as a biomarker for monitoring disease recurrence in IBD patients and may be a strong marker of disease activity in UC patients, potentially outperforming CRP. Clinically, the identification of a divergence of biological age from chronological age, or the presence of a negative aging trajectory, may highlight IBD patients at greatest risk of disease progression, allowing early therapeutic intervention, including medicines that directly modulate aging processes.

To the best of our knowledge, this is the first study to report on the third generation DNA methylation biomarker of pace of aging (DunedinPACE - for Pace of Aging Calculated from the Epigenome), in addition to a range of second-generation ‘epigenetic’ clocks, and their association with IBD, including disease recurrence, activity, subtypes and other relevant clinicopathological variables.

## List of abbreviations

IBD: Inflammatory Bowel Diseases
UC: Ulcerative Colitis
CD: Crohn’s Disease
DNAm: DNA Methylation
GEO: Gene Expression Omnibus
DMP: Differentially Methylated Positions
BH: Benjamini Hochberg
CRP: C-reactive Protein
ROC: Receiver Operator Characteristic
AUC: Area under the ROC curve
PBMC: Peripheral Blood Mononuclear Cell
RISK: Risk Stratification and Identification of Immunogenetic and Microbial Markers of Rapid Disease Progression in Children with Crohn’s Disease.

## Availability of data and materials

The publicly available datasets analysed during the current study are available in the Gene Expression Omnibus repository and are accessible through GEO series accessions GSE87648 and GSE112611. Data supporting some of the findings of this study are not openly available due to reasons of sensitivity and are available from the corresponding author upon reasonable request.

## Authors’ contributions

TMM conceived the study and supervised the project. TD performed all statistical and DNA methylation analysis. TD, SJD and TMM contributed to the computational interpretation of the results and were involved in the overall manuscript conceptualisation. TMM, EMcD and HM contributed clinical data and provided DNA methylation data for the study. EMcD and HM conceptualised data collection protocols and created variables for the Dublin IBD cohort. TD and TMM drafted the manuscript. All authors reviewed and approved the final manuscript.

## Acknowledgements

We would like to thank the Irish Centre for High End Computing (https://www.ichec.ie/) and the MeluXina platform (https://www.luxprovide.lu/meluxina/) for the use of their HPC infrastructure and support, and the School of Computer Science (Technological University Dublin) for provision of computing cluster resources.

## Ethics declarations

### Ethics approval and consent to participate

Ethical approval was obtained from St. Vincent’s University Hospital (Dublin) Research and Ethics Committee.

### Consent for publication

Not applicable.

