## Supplementary Information for "Analysis of blood-based DNA Methylation signatures of aging and disease progression in IBD patients"

Table S1: Top ranked differentially methylated positions in IBD long recurrence (n=54) compared with non-recurrence (n=92). Adjusted p-value is BH-adjusted.

| **Probe ID (CpG)** | **P-value** | **Effect Size (regression coefficient)** | **Adjusted P-value** | **Hg19** | **Illumina Annotation** | **Probe Type** |
| --- | --- | --- | --- | --- | --- | --- |
| **cg03583111** | **4.96E-08** | **0.015** | **0.021** | Chr2:  220283442 | *DES* | II |
| cg13036855 | 4.32E-07 | 0.013 | 0.093 | Chr3:  53880717 | *CHDH;*  *IL17RB* | I |
| cg16359901 | 1.22E-06 | 0.012 | 0.149 | Chr1: 6269635 | *RNF207* | II |
| cg09816693 | 1.60E-06 | 0.015 | 0.149 | Chr14: 74706113 | *VSX2* | II |
| cg10634273 | 1.73E-06 | -0.013 | 0.149 | Chr1:  84864163 | *DNASE2B; UOX* | II |
| cg14638662 | 2.96E-06 | 0.019 | 0.185 | Chr17:  73033177 |  | II |
| cg22876092 | 3.09E-06 | -0.020 | 0.185 | Chr10:  88126287 | *GRID1* | I |
| cg12789522 | 3.43E-06 | -0.011 | 0.185 | Chr15: 78423758 | *CIB2* | I |
| cg24897126 | 4.62E-06 | 0.022 | 0.206 | Chr1:  6474383 |  | II |
| cg03907847 | 4.78E-06 | 0.012 | 0.206 | Chr13:  28495908 | *PDX1* | II |

Table S2: Top ranked differentially methylated positions in IBD short recurrence (n=78) compared with non-recurrence (n=68). Adjusted p-value is BH-adjusted.

| **Probe ID (CpG)** | **P-value** | **Effect Size (regression coefficient)** | **Adjusted P-value** | **Hg19** | **Illumina Annotation** | **Probe Type** |
| --- | --- | --- | --- | --- | --- | --- |
| cg26010908 | 2.86E-06 | 0.003 | 0.731 | Chr6:  31628367 | *C6orf47* | II |
| cg08907118 | 3.39E-06 | -0.012 | 0.731 | Chr16:  27482516 | *GTF3C1* | II |
| cg23124486 | 1.26E-05 | 0.011 | 0.990 | Chr6:  33166165 | *RXRB* | II |
| cg23866916 | 2.43E-05 | -0.014 | 0.990 | Chr19:  1155738 | *SBNO2* | II |
| cg08080060 | 2.90E-05 | 0.005 | 0.990 | Chr19:  41307090 | *EGLN2* | I |
| cg06507434 | 3.08E-05 | 0.004 | 0.990 | Chr11:  64597280 | *CDC42BPG* | I |
| cg07034103 | 4.65E-05 | -0.011 | 0.990 | Chr13:  39885248 |  | II |
| cg04757002 | 4.72E-05 | 0.007 | 0.990 | Chr2;  20424804 | *SDC1* | I |
| cg19279042 | 4.97E-05 | 0.011 | 0.990 | Chr6:  31550090 | *LTB* | II |
| cg11229610 | 5.66E-05 | 0.008 | 0.990 | Chr12:  49412926 | *MLL2;*  *PRKAG1* | II |

Table S3: Association of standardised DNAm aging signatures with IBD vs. control status for the unadjusted and adjusted (sex, smoking, blood cell concentrations) logistic regression models.

|  | **IBD vs. Control**  **(Not covariate adjusted)** | | | **IBD vs. control**  **(Covariate Adjusted)** | | |
| --- | --- | --- | --- | --- | --- | --- |
| **DNAm Signature** | **P-value** | **Adjusted P-value** | **OR** | **P-value** | **Adjusted P-value** | **OR** |
| DunedinPACE | **0.003** | **0.012** | 1.876 | **0.024** | 0.056 | 3.628 |
| AgeAccelGrim | **0.004** | **0.012** | 1.836 | **0.037** | 0.065 | 2.692 |
| AgeAccelGrim2 | **0.008** | **0.019** | 1.699 | **0.017** | 0.056 | 3.724 |
| Epigenetic Age (Zhang)AdjAge | 0.334 | 0.560 | 0.838 | 0.154 | 0.154 | 1.943 |
| AgeAccelPheno | 0.445 | 0.560 | 1.149 | 0.063 | 0.088 | 2.508 |
| MI-EN TLAdjAge | 0.480 | 0.560 | 1.138 | 0.077 | 0.090 | 0.450 |
| DNAmTLAdjAge | 0.812 | 0.812 | 0.958 | **0.009** | 0.056 | 0.299 |

Table S4: Association of standardised DNAm aging signatures with CD vs. control status for the unadjusted and adjusted (sex, smoking, blood cell concentrations) logistic regression models.

|  | **CD vs. Control**  **(Not covariate adjusted)** | | | **CD vs. control**  **(Covariate Adjusted)** | | |
| --- | --- | --- | --- | --- | --- | --- |
| **DNAm Signature** | **P-value** | **Adjusted P-value** | **OR** | **P-value** | **Adjusted P-value** | **OR** |
| DunedinPACE | **0.0006** | **0.0022** | 2.2741 | **0.006** | **0.029** | 7.003 |
| AgeAccelGrim | **0.0006** | **0.0022** | 2.2246 | **0.012** | **0.029** | 4.448 |
| AgeAccelGrim2 | **0.0019** | **0.0043** | 2.0097 | **0.010** | **0.029** | 6.426 |
| AgeAccelPheno | 0.2009 | 0.3516 | 1.2860 | 0.056 | 0.066 | 3.242 |
| DNAmTLAdjAge | 0.5712 | 0.7997 | 0.8960 | **0.021** | **0.038** | 0.266 |
| Epigenetic Age (Zhang)AdjAge | 0.7389 | 0.8621 | 0.9372 | 0.117 | 0.117 | 2.408 |
| MI-EN TLAdjAge | 0.8856 | 0.8856 | 0.9724 | **0.041** | 0.057 | 0.334 |

Table S5: Association of standardised DNAm aging signatures with UC vs. control status for the unadjusted and adjusted (sex, smoking, blood cell concentrations) logistic regression models.

|  | **UC vs. Control**  **(Not covariate adjusted)** | | | **UC vs. control**  **(Covariate Adjusted)** | | |
| --- | --- | --- | --- | --- | --- | --- |
| **DNAm Signature** | **P-value** | **Adjusted P-value** | **OR** | **P-value** | **Adjusted P-value** | **OR** |
| Epigenetic Age (Zhang)AdjAge | 0.060 | 0.235 | 0.657 | 0.820 | 0.82 | 1.124 |
| DunedinPACE | 0.112 | 0.235 | 1.429 | 0.244 | 0.427 | 2.148 |
| AgeAccelGrim | 0.131 | 0.235 | 1.394 | 0.429 | 0.580 | 1.617 |
| MI-EN TLAdjAge | 0.147 | 0.235 | 1.365 | 0.497 | 0.580 | 0.685 |
| AgeAccelGrim2 | 0.168 | 0.235 | 1.348 | 0.175 | 0.427 | 2.437 |
| DNAmTLAdjAge | 0.693 | 0.729 | 1.086 | **0.038** | 0.264 | 0.298 |
| AgeAccelPheno | 0.729 | 0.729 | 0.930 | 0.196 | 0.427 | 1.932 |

Table S6: Association of standardised DNA methylation aging signatures with IBD/control status for both unadjusted and adjusted (sex, blood cell concentrations) logistic regression models for the GSE87648 replication data set.

|  | **IBD vs. Control**  **(Not covariate adjusted)** | | | **IBD vs. control**  **(Covariate Adjusted)** | | |
| --- | --- | --- | --- | --- | --- | --- |
| **DNAm Signature** | **P-value** | **Adjusted P-value** | **OR** | **P-value** | **Adjusted P-value** | **OR** |
| DunedinPACE | **8.078e-21** | **2.423e-20** | 6.308 | **1.782e-15** | **5.346e-15** | 5.707 |
| AgeAccelGrim2 | **1.109e-15** | **1.663e-15** | 3.451 | **4.147e-09** | **6.220e-09** | 2.708 |
| AgeAccelGrim | **2.716e-08** | **2.716e-08** | 1.961 | **9.836e-05** | **9.836e-05** | 1.806 |

Table S7: Association of standardised DNA methylation aging signatures with CD/control status for both unadjusted and adjusted (sex, blood cell concentrations) logistic regression models for the GSE87648 replication data set.

|  | **CD vs. Control**  **(Not covariate adjusted)** | | | **CD vs. control**  **(Covariate Adjusted)** | | |
| --- | --- | --- | --- | --- | --- | --- |
| **DNAm Signature** | **P-value** | **Adjusted P-value** | **OR** | **P-value** | **Adjusted P-value** | **OR** |
| DunedinPACE | **7.008e-17** | **2.102e-16** | 8.217 | **5.634e-13** | **1.690e-12** | 7.238 |
| AgeAccelGrim2 | **2.229e-14** | **3.343e-14** | 4.594 | **3.192e-10** | **4.788e-10** | 3.894 |
| AgeAccelGrim | **6.646e-09** | **6.646e-09** | 2.365 | **1.566e-06** | **1.566e-06** | 2.484 |

Table S8: Association of standardised DNA methylation aging signatures with UC/control status for both unadjusted and adjusted (sex, blood cell concentrations) logistic regression models for the GSE87648 replication data set.

|  | **UC vs. Control**  **(Not covariate adjusted)** | | | **UC vs. control**  **(Covariate Adjusted)** | | |
| --- | --- | --- | --- | --- | --- | --- |
| **DNAm Signature** | **P-value** | **Adjusted P-value** | **OR** | **P-value** | **Adjusted P-value** | **OR** |
| DunedinPACE | **5.753e-14** | **1.726e-13** | 4.511 | **4.644e-10** | **1.393e-09** | 4.330 |
| AgeAccelGrim2 | **8.908e-09** | **1.336e-08** | 2.479 | **2.117e-04** | **3.175e-04** | 1.942 |
| AgeAccelGrim | **5.544e-04** | **5.544e-04** | 1.578 | **4.930e-02** | **4.930e-02** | 1.388 |

Table S9: Association of standardised epigenetic age acceleration (z-scores) measures with disease trajectory (short recurrence vs. non-recurrence and long recurrence vs. non-recurrence) in the unadjusted logistic regression model. Significant values are shown in bold.

|  | **Short Recurrence** | | | **Long Recurrence** | | |
| --- | --- | --- | --- | --- | --- | --- |
| **DNAm Signature** | **P-value** | **Adjusted P-value** | **OR** | **P-value** | **Adjusted P-value** | **OR** |
| DunedinPACE | **0.032** | 0.222 | 1.454 | 0.986 | 0.986 | 0.997 |
| AgeAccelGrim | 0.192 | 0.474 | 1.245 | 0.182 | 0.854 | 1.264 |
| AgeAccelGrim2 | 0.203 | 0.474 | 1.239 | 0.457 | 0. 854 | 1.137 |
| AgeAccelPheno | 0.741 | 0.983 | 1.057 | 0.746 | 0.871 | 1.057 |
| DNAmTLAdjAge | 0.850 | 0.983 | 1.032 | 0.272 | 0. 854 | 1.210 |
| MI-EN TLAdjAge | 0.962 | 0.983 | 0.992 | 0.488 | 0. 854 | 0.888 |
| Epigenetic Age (Zhang)AdjAge | 0.983 | 0.983 | 0.996 | 0.613 | 0.858 | 0.917 |

Table S10: Association of standardised epigenetic age acceleration measures with disease trajectory (short recurrence vs. non-recurrence and long recurrence vs. non-recurrence) in the adjusted multivariable logistic regression model (sex, smoking, blood cell concentrations).

|  | **Short Recurrence** | | | **Long Recurrence** | | |
| --- | --- | --- | --- | --- | --- | --- |
| **DNAm Signature** | **P-value** | **Adjusted P-value** | **OR** | **P-value** | **Adjusted P-value** | **OR** |
| DunedinPACE | **0.036** | 0.249 | 1.733 | 0.739 | 0.862 | 0.920 |
| AgeAccelGrim2 | 0.214 | 0.523 | 1.424 | 0.185 | 0.483 | 1.471 |
| AgeAccelGrim | 0.224 | 0.523 | 1.380 | 0.071 | 0.483 | 1.664 |
| MI-EN TLAdjAge | 0.508 | 0.882 | 0.864 | 0.236 | 0.483 | 0.769 |
| DNAmTLAdjAge | 0.720 | 0.882 | 1.094 | 0.355 | 0.496 | 1.264 |
| Epigenetic Age (Zhang)AdjAge | 0.756 | 0.882 | 1.067 | 0.928 | 0.928 | 0.981 |
| AgeAccelPheno | 0.912 | 0.912 | 1.029 | 0.276 | 0.483 | 1.327 |

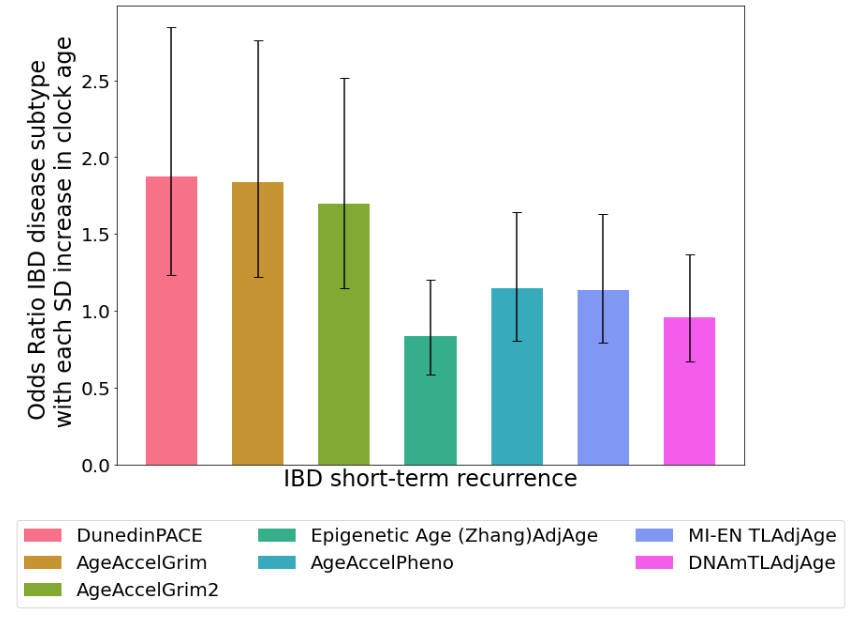

Figure S1: Odds ratios of DNAm signatures of aging in blood for IBD vs. control status (n=184). Error bars indicate 95% confidence intervals.

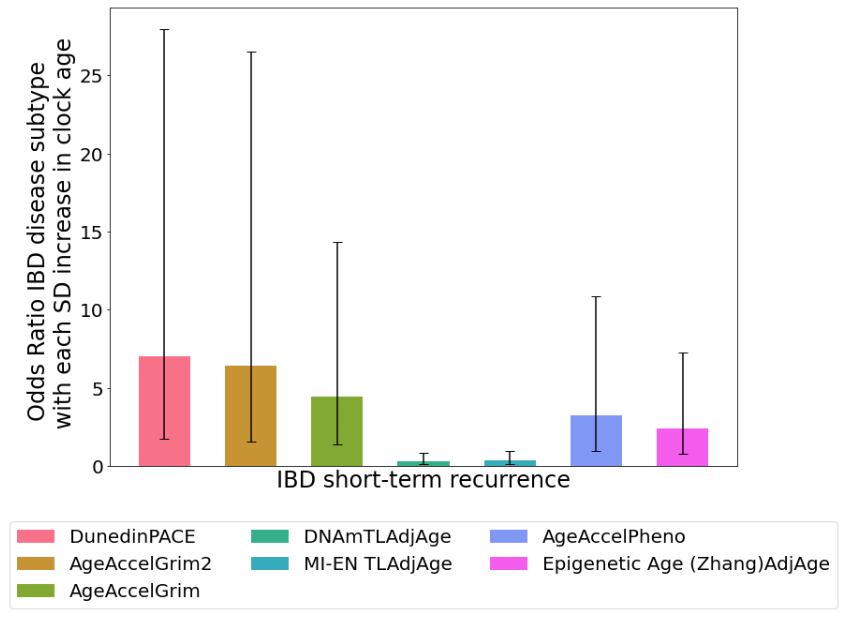

Figure S2: Odds ratios of DNAm signatures of aging in blood for CD vs. control status. Odds ratios adjusted for sex, smoking status and blood cell concentrations (n=125). Error bars indicate 95% confidence intervals.

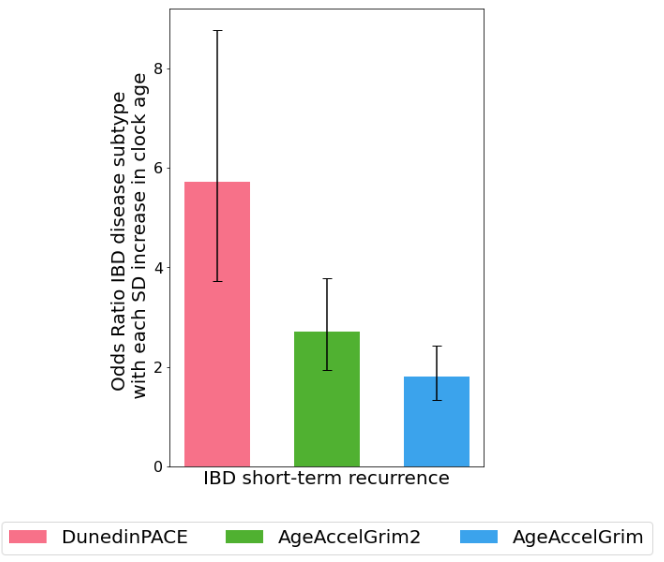

Figure S3: Odds ratios of DNAm signatures of aging in blood for IBD vs. control status. Odds ratios adjusted for sex and blood cell concentrations (n=379). Error bars indicate 95% confidence intervals.

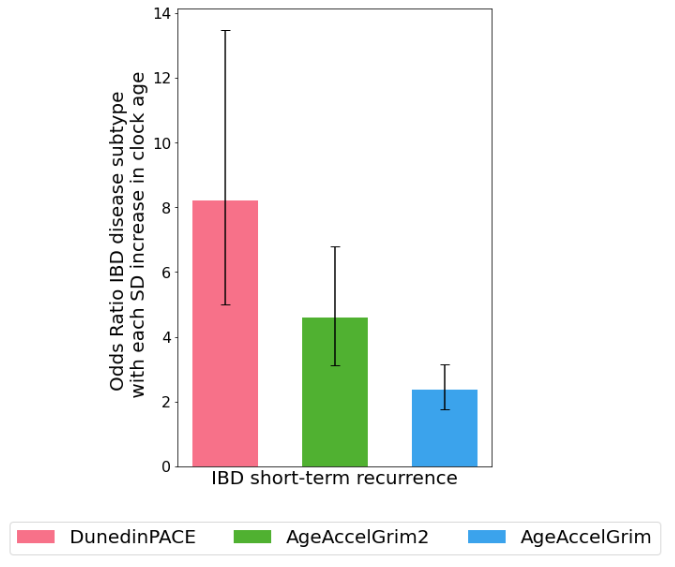

Figure S4: Odds ratios of DNAm signatures of aging in blood for CD/control status (n=278). Error bars indicate 95% confidence intervals (model without covariate adjustment).

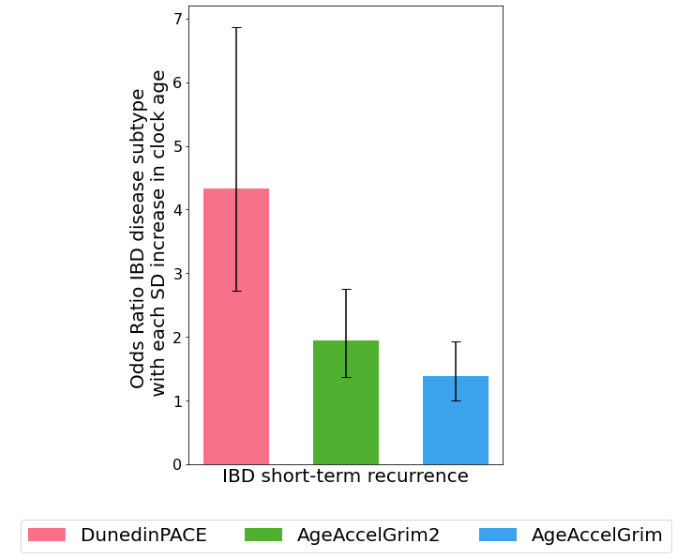

Figure S5: Odds ratios of DNAm signatures of aging in blood for UC/control status (n=276). Error bars indicate 95% confidence intervals (model with covariate adjustment).

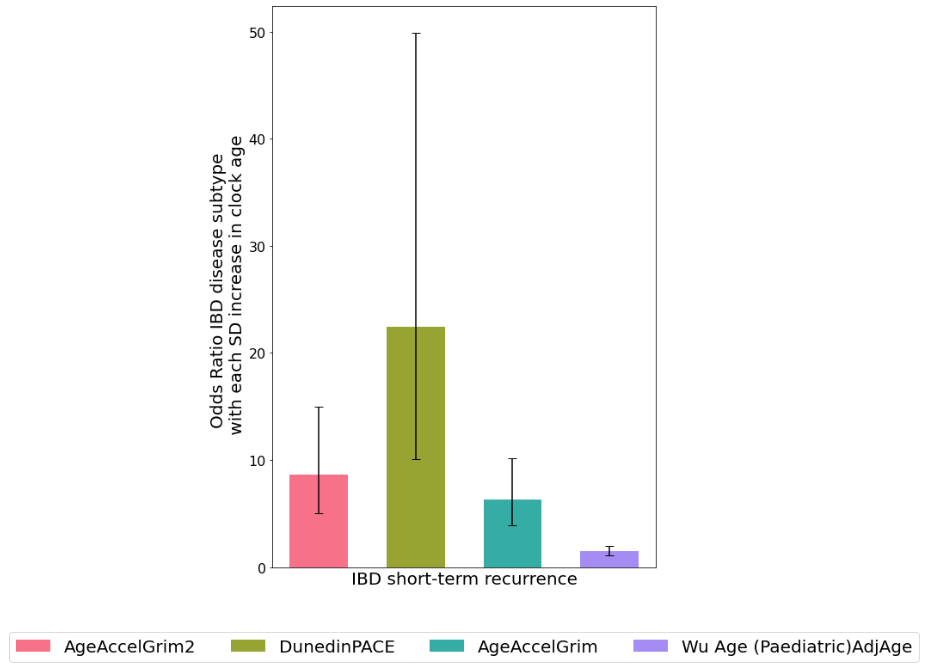

Figure S6: Odds ratios of DNAm signatures of aging in blood for CD/control status (n=238). Error bars indicate 95% confidence intervals (model without covariate adjustment).

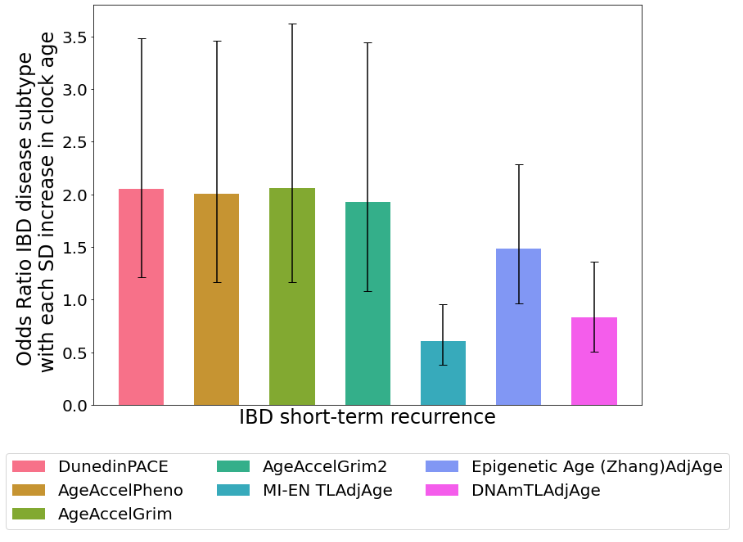

Figure S7: Odds ratios of DNAm signatures of aging in blood for IBD disease subtype (UC vs. CD). Odds ratios adjusted for sex, smoking status and white blood cell concentrations (n=146). Error bars indicate 95% confidence intervals.

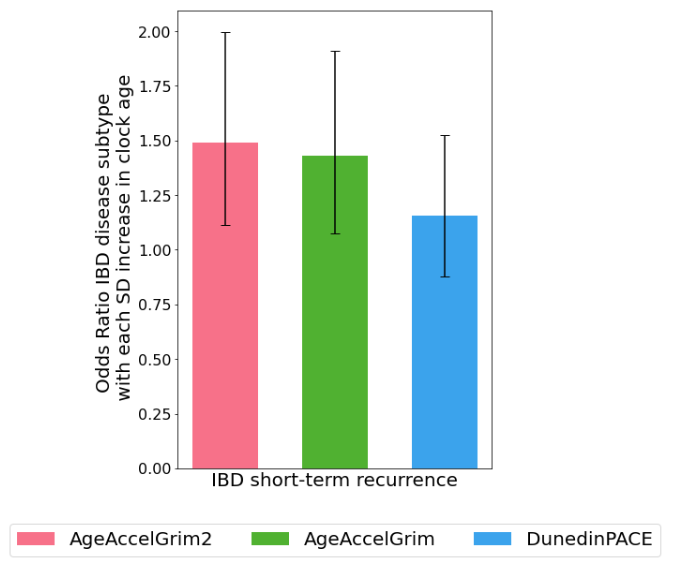

Figure S8: Odds ratios of DNAm signatures of aging in blood for IBD disease subtype (UC vs. CD) (n=204). Error bars indicate 95% confidence intervals (model without covariate adjustment).
